# Structural mechanisms of drebrin-mediated F-actin network modulation

**DOI:** 10.64898/2025.12.01.691618

**Authors:** W Zhao, G Abis, F Oozeer, T Mulvaney, N Nagar, M Topf, PR Gordon-Weeks, MR Conte, J Atherton

**Affiliations:** Randall Centre for Cell and Molecular Biophysics, King’s College London – New Hunt’s House, Guy’s Campus, London SE1 1UL, United Kingdom; Centre for Developmental Neurobiology, King’s College London – New Hunt’s House, Guy’s Campus, London SE1 1UL, United Kingdom; Centre for Structural Systems Biology (CSSB), Hamburg, Germany; Department of Integrative Virology, Leibniz Institute of Virology (LIV), Hamburg, Germany.; Institute for Molecular Virology and Tumour virology, University Medical Center Hamburg-Eppendorf (UKE), Hamburg, Germany

## Abstract

Drebrin modulates F-actin networks and links them to other intracellular components, regulating crucial processes including neuritogenesis, synaptic plasticity, virus internalisation and cancer invasion. Using single-particle cryo-EM we characterise drebrin’s interaction with F-actin through two separate conserved actin binding domains (ABD1 and ABD2), revealing structural bases for its F-actin-modulating properties. We describe a multimodal interaction where drebrin’s ABD1 can adopt two conformations and a long flexible loop connecting to ABD2 allows the two ABDs to occupy multiple relative positions along F-actin. Despite harbouring two separated ABDs, we find that drebrin is not a strong direct F-actin bundler. Drebrin’s ABDs bind across multiple actin protomers and their subdomains and modify the longitudinal inter-protomer interface, explaining their F-actin stabilising properties. Furthermore, we show drebrin’s binding site on F-actin is shared with other critical actin-binding and regulatory proteins, explaining their competitive displacement.

## Introduction

Filamentous actin (F-actin) dynamically adopts various higher-order architectures to perform defined sub-cellular functions. Numerous actin-binding proteins (ABPs), which often compete with one another or act cooperatively, modulate the growth and stability of individual filaments or organise multiple filaments into various bundled and branched networks.

Drebrin, a vertebrate actin-depolymerizing factor homology (ADFH) protein family member, contains an N-terminal ADFH domain, followed by either one or several F-actin binding regions of predicted helical content^1^. The majority of studies find the ADFH domain does not bind F-actin^1,2^, unlike for other ADFH protein family members such as cofilin and actin-binding protein 1 (ABP1)^3^. Drebrin’s two isoforms, drebrin-A and drebrin-E, differ in the inclusion or exclusion respectively of a 45 amino acid insert following the F-actin binding region. While drebrin-A is expressed in adult neurons only, drebrin-E is expressed in developing neurons and multiple other cell types.

Drebrin has key functions in specific cellular subdomains. In developing neurons, drebrin-E regulates neuronal migration, process outgrowth and filopodial stability, locating predominantly to growth cone transition zone F-actin arcs and peripheral zone filopodia bases^4–12^. In both neuronal and non-neuronal cell lines, drebrin overexpression strongly induces filopodia formation^11–14^. In mature neurons, drebrin-A is concentrated on dendritic spine F-actin, functioning in synapse formation and remodelling and learning and memory^15–22^. Postsynaptic drebrin expression abnormalities are found in epilepsy^23^, Down syndrome and Alzheimer’s disease (AD)^24–27^, with evidence from AD mouse models suggesting drebrin is a modulator of disease progression^28,29^. Interestingly, neuronal drebrin is also a modulator of opioid addiction^30^.

Drebrin structurally supports other cell-cell junctions such as gap junctions^31–33^ and adherens junctions^34^ through linking F-actin at the recipient side with key structural and signalling proteins. Drebrin recruits CXCR4 to T-cell F-actin at the immune synapse during the immune response^35^ and negatively regulates HIV entry through this pathway^36^. Drebrin’s modulation of endocytotic F-actin networks also affects rotavirus and pseudorabies virus entry^37,38^. In the developing lens, F-actin localised drebrin is required for its proper macro-organisation^39^. Drebrin reduces vascular smooth muscle cell activation, migration and proliferation through F-actin interactions, attenuating atherosclerosis ^40,41^. Drebrin is expressed in myofibroblasts in fibrotic heart, lung and liver where it exerts profibrotic affects ^42,43^. Finally, drebrin upregulation and F-actin interactions are implicated in the metastasis and invasion of various cancers^44–50^.

Drebrin’s functions are critically linked to F-actin interactions through several potential molecular mechanisms. Firstly, drebrin exerts direct effects on F-actin stability and architecture. For example, drebrin has been reported to bind cooperatively, directly inhibit F-actin depolymerisation, increase filament stiffness, reduce filament curvature and remodel F-actin by increasing the cross-over repeat from ∼36 nm to ∼40 nm^51–55^. While one study suggested drebrin can bundle F-actin directly^12^, others have found no direct bundling effect^55–58^. Secondly, drebrin competitively displaces other ABPs from F-actin, with downstream consequences. To date, drebrin has been demonstrated to compete with F-actin depolymerase cofilin, F-actin-bundlers fascin and α-actinin, tropomyosin and myosins, suppressing their activities^55,59–62^. Finally, drebrin links the F-actin cytoskeleton to an increasing array of structural and signalling proteins^1^. For example, drebrin links F-actin to microtubules through binding EB3, encouraging microtubule entry/retention in filopodia^7,11^ and promoting prostate cancer cell invasion^63^.

Despite the centrality of drebrin’s interaction with and regulation of F-actin in driving its cellular functions, a structural understanding is lacking. Here, we present ∼2.3 Å resolution cryo-electron microscopy (cryo-EM) reconstructions of drebrin’s actin-binding region bound to F-actin, revealing two conserved actin-binding domains (ABDs). The two ABDs compete sterically to define drebrin stoichiometry and each bind across two actin protomers and multiple F-actin subdomains (SDs) to stabilise the filament. These ABDs also displace various ABPs via steric competition for F-actin binding sites. The first ABD adopts two conformations upon F-actin, with implications for binding stoichiometry and ABP competition. Despite being separated by a flexible region, the two ABDs do not visibly bundle F-actin. Furthermore, no significant drebrin-induced remodelling of F-actin is observed.

## Results

### Drebrin binds F-actin via two actin-binding domains

We first purified drebrin-E^135–355^, thought to contain the core F-actin binding region, and a longer drebrin-E^1–431^ construct that also contains the globular ADFH domain which interacts with actin in other ADFH protein family members^3^ and the PP domain (predicted unstructured) (Fig. 1a). While circular dichroism (CD) indicated that both constructs are chiefly composed of random coil and α-helical structure, drebrin-E^1–431^ had a slight increase in β-sheet content, consistent with the presence of a folded ADFH domain (Extended Data Fig. 1a). Structural prediction of drebrin-E with AlphaFold3^64^ and JPred4^65^ suggested that the F-actin binding region within amino acids 135-355 is composed of predominantly α-helical content with some random coil (Extended Data Fig. 1b and c).

**Figure 1.**
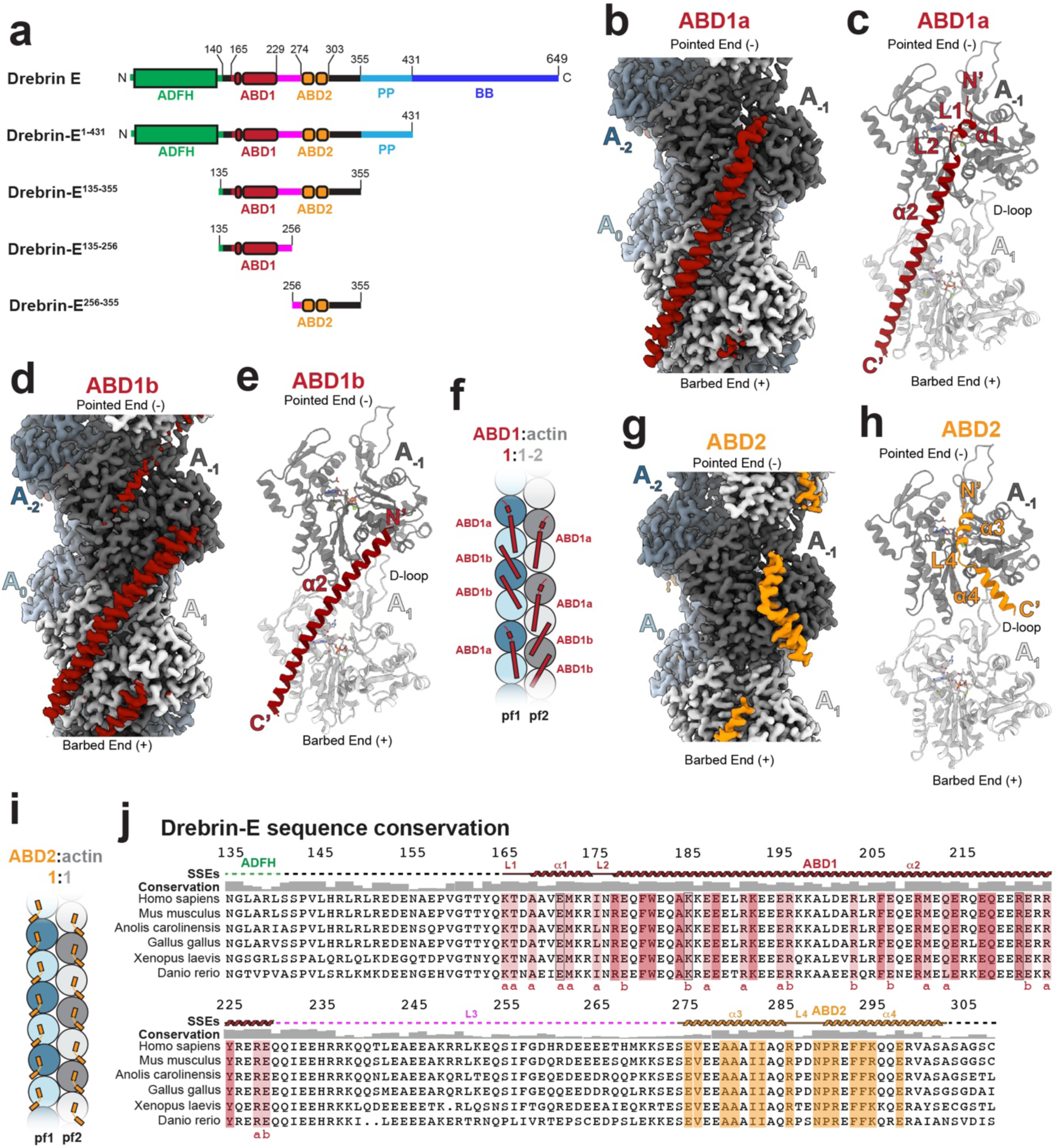
Drebrin’s ABDs each bind across two F-actin protomers, with ABD1 adopting two conformations. a) Domain diagram for human drebrin-E and truncated drebrin-E constructs used in this study. b) Cryo-EM reconstruction (from our drebrin-E^135–355^ dataset) and c) corresponding atomic model (below) for the ABD1a conformation. d) Cryo-EM reconstruction (from our drebrin-E^135–355^ dataset) and e) corresponding atomic model (below) for the ABD1b conformation. f) Schematic model showing ABD1 assuming both ABD1a and ABD1b conformations on F-actin, such that the stoichiometry of ABD1 alone varies between 1:1-2 for F-actin. g) Cryo-EM reconstruction (from our drebrin-E^135–355^ dataset) and h) corresponding atomic model for ABD2. i) Schematic model showing ABD2 alone assuming a single conformation with 1:1 stoichiometry with actin. j) A multiple sequence alignment of human drebrin-E residues 135-308 with canonical drebrin orthologue sequences from 113 other vertebrate species (acquired from the Ensembl database^113^) was performed with the MUSCLE algorithm^114^. The *Homo sapiens* sequence alongside a reduced set of 5 vertebrate class model species (mammals, reptiles, birds, amphibians and fish) is shown, with *Homo sapiens* sequence numbering, secondary structure elements (SSEs) identified by cryo-EM and sequence conservation from the larger 113 species alignment shown above. F-actin contacting residues are highlighted in red and orange for ABD1 and ABD2 respectively. Those contacting F-actin in both ABD1 conformations are highlighted in dark red, those in one or the other conformation are highlighted in light red, with the conformation indicated below (a= ABD1a, b=ABD1b). Residues within black boxes were mutated for the analysis presented in Extended Data Fig. 7. In panels f and i pf1 and 2 = protofilaments 1 and 2 of F-actin. In panels b-e and g-h F-actin protomers are labelled from A_-2_ to A_1_ (from pointed end to barbed end).

We then collected cryo-EM datasets of both drebrin-E^135–355^ and drebrin-E^1–431^-decorated F-actin for single-particle analysis. Initial processing revealed additional density for drebrin-E on F-actin that was indistinguishable for our drebrin-E^135–355^ and drebrin-E^1–431^ datasets (Extended Data Fig. 1d and e), confirming that the ADFH or PP domains do not constitute additional F-actin binding regions. The density was of poorer quality than expected and much was only apparent at less conservative density thresholds. Considering drebrin contains two reported ABDs^12^, we reasoned that if each ABD occupies exclusive actin protomers along the filament due to shared binding sites, their densities would likely be aberrantly averaged during image processing due to F-actin density dominating image alignment. This was supported by structural prediction of the drebrin-E^135–355^-F-actin complex with AlphaFold3^64^, which suggested that multiple sections of the F-actin binding region share binding surfaces on F-actin (Extended Data Fig. 1f). To overcome this, we collected cryo-EM datasets of F-actin decorated with either drebrin-E^135–256^ or drebrin-E^256–355^ constructs which each contain only one of two reported ABDs that have independent actin-binding activities^12^ (Fig. 1a and Table 1). Picked particles corresponding to decorated segments of F-actin filaments were subjected to image processing including focussed classification such as to identify different conformations of drebrin-E^135–256^ or drebrin-E^256–355^ if present (Extended Data Fig. 2).

**Table 1.** Cryo-EM data collection, refinement and validation statistics.

|  | Drebrin-E<sup>135-355</sup> ABD1a<br>F-actin (EMD-55893,<br>PDB: 9TG8) | Drebrin-E<sup>135-355</sup> ABD1b<br>F-actin (EMD-55894,<br>PDB: 9TG9) | Drebrin-E<sup>135-355</sup> ABD2 F-actin (EMD-55895,<br>PDB: 9TGA) |
| --- | --- | --- | --- |
| **Data collection and processing** |  |  |  |
| Magnification (actual) | 150,376 | 150,376 | 150,376 |
| Voltage (kV) | 300kv | 300kv | 300kv |
| Electron exposure (e<sup>-</sup><br>/Å<sup>2</sup>) | 45 | 45 | 45 |
| Defocus range (μm) | -0.7 to -2.5μm | -0.7 to -2.5μm | -0.7 to -2.5μm |
| Pixel size (Å) | 0.931 | 0.931 | 0.931 |
| Symmetry imposed | C1 | C1 | C1 |
| Initial micrographs (#) | 3243 | 3243 | 3243 |
| Final single particles<br>used (#) | 102,155 | 115,898 | 149,259 |
| Map resolution (Å) | 2.37 | 2.34 | 2.29 |
| FSC threshold | 0.143 | 0.143 | 0.143 |
| Map resolution range<br>(Å) | 2.36-3.33 | 2.30-3.22 | 2.16-3.00 |
| Measured helical<br>symmetry |  |  |  |
| Helical rise (Å) | 27.6 | 27.6 | 27.6 |
| Helical twist (°) | -166.653 | -166.653 | -166.653 |
| **Refinement** |  |  |  |
| Initial model used | PDB: 8a2s (actin) | PDB: 8a2s (actin) | PDB: 8a2s (actin) |
| Model resolution (Å) | 2.43 | 2.31 | 2.23 |
| FSC threshold | 0.5 | 0.5 | 0.5 |
| Map sharpening B<br>factor (Å<sup>2</sup>) | -32.29 | -30.28 | -30.87 |
| Model composition |  |  |  |
| Non-hydrogen atoms | 6701 | 6534 | 6432 |
| Protein residues | 805 | 794 | 770 |
| Waters | 265 | 184 | 352 |
| Ligands | 4 | 4 | 4 |
| B factors (Å<sup>2</sup>) |  |  |  |
| Protein | 44.26 | 43.54 | 34.26 |
| Waters | 41.93 | 41.36 | 34.05 |
| Ligand | 36.01 | 33.09 | 24.78 |
| R.m.s. deviations |  |  |  |
| Bond lengths (Å) | 0.003 | 0.004 | 0.004 |
| Bond angles (°) | 0.573 | 0.965 | 1.009 |
| Validation |  |  |  |
| MolProbity score | 1.16 | 1.00 | 1.15 |
| Clashscore | 3.76 | 2.22 | 3.57 |
| Poor rotamers (%) | 0.15 | 0.44 | 0.15 |
| Ramachandran plot |  |  |  |
| Favored (%) | 98.99 | 99.23 | 99.08 |
| Allowed (%) | 1.01 | 0.77 | 0.92 |
| Disallowed (%) | 0.00 | 0.00 | 0.00 |
| Occupancy |  |  |  |
| Mean | 0.76 | 0.79 | 0.77 |
| Occ = 1 (%) | 1.01 | 0.40 | 0.51 |
| 0 < Occ < 1(%) | 98.96 | 99.56 | 99.49 |
| Occ > 1(%) | 0.00 | 0.00 | 0.00 |

This strategy revealed that drebrin-E^135–256^ and drebrin-E^256–355^ each contain an ABD domain which we refer to as ABD1 and ABD2 respectively, with ABD1 adopting two alternative conformations (Extended Data Fig. 2). Densities for ABD1 and ABD2 were found at overlapping sites on F-actin such that aberrant averaging of the two domains likely occurred during attempts to image process datasets of F-actin decorated with longer drebrin-E^135–355^ and drebrin-E^1–431^constructs (Extended Data Fig. 1d and e). These three conformations (two for ABD1 and one for ABD2) were then used as references to guide a focussed supervised classification strategy to identify sites occupied with these ABD1 or ABD2 conformations within the drebrin-E^135–355^-decorated F-actin dataset (Extended Data Fig. 3). The resultant ∼2.3 Å resolution cryo-EM maps were of sufficient quality to build atomic models of the interface of F-actin with ordered drebrin ABD1 and ABD2 sequences (Fig. 1b-j, Extended Data Fig. 4a-I and Table 1).

### Each drebrin ABD binds two F-actin protomers

Focussed classification identified two distinct conformations of ABD1 with roughly equal occupancy and partially-overlapping F-actin binding surfaces which we will refer to as ABD1a and ABD1b (Fig. 1b-e, Extended Data Fig. 2-4). In both conformations, a long α-helix (amino acids R177-E227, helix-α2) straddles two protomers along a single F-actin protofilament (A_-1_ and A_1_ moving in the pointed to barbed end direction) but adopts different trajectories, increasingly diverging laterally from one another towards their N-termini on A^-1^ (Fig. 1b and e). Furthermore, in ABD1a K165-N176 forms an additional ordered F-actin binding region (associated with A^-1^) containing a short α-helix (helix-α1) which is connected to the proceeding long α2 by a short loop (L2) (Fig. 1c and j). ABD1 map resolution decreases and model B-factor increases towards the helix-α2 C-terminus, suggesting it has somewhat higher mobility (Extended Data Fig. 4d-f).

ABD1a has proximal binding surfaces for its N and C-terminal contacts on each substrate protomer, therefore due to steric restraints a single ABD1a exclusively occupies a single protomer pair along the protofilament (Fig. 1c and f, Extended Data Fig. 5a). In contrast, ABD1b’s modified trajectory shifts its binding surface on A_-1_ such that binding of another ABD1’s C-terminal end is now sterically permitted (Fig. 1e and f, Extended Data Fig. 5b and c). Therefore, adoption of a proportion of the ABD1b conformation permits a higher potential occupancy of ABD1 on F-actin (Fig. 1f).

In contrast, only one ABD2 conformation was found, corresponding to drebrin-E amino acids 274-303 (Fig. 1g and h). ABD2 is composed of two α-helices (helix-α3 and helix-α4) joined by a short, ordered loop (L4) (Fig. 1h). ABD2 primarily contacts a single actin protomer (A_-1_), however the C-terminal portion of helix-α4 also contacts the D-loop of the next protomer along the protofilament (A_1_, Fig. 1h). Although ABD2 binds across two actin protomers, it binds to F-actin with a stoichiometry of 1:1 (Fig. 1i).

### F-actin-interacting residues in drebrin ABDs are highly conserved and important for drebrin function

To identify functionally important residues, we integrated our structural data with drebrin sequence conservation across vertebrate species and extracted predicted substitution pathogenicity scores from AlphaMissense^66^. High conservation was found in ABD1 and ABD2, particularly in F-actin-interacting residues (Fig. 1j and Extended Data Fig. 6a-e). High conservation was found in the L1-helix-α1-L2 sequence and downstream residues which exclusively interact with F-actin in ABD1a (Fig. 1h and Extended Data Fig. 6c). However, several residues specifically interacting with F-actin in only ABD1b were also found to be highly conserved (Fig. 1h and Extended Data Fig. 6d). Taken together, these results highlight the conserved functional importance of F-actin association via both ABDs and further suggest that both ABD1 conformations serve conserved physiological functions.

Transfection of an eYFP-tagged drebrin-E^135–256^ construct^12^ (containing ABD1 alone) into COS7 cells induces filopodia when they are usually rare^11,12^. We tested the functional importance of each ABD1 conformation for filopodial induction by expressing this construct with substitutions targeted to conserved charged residues that clearly either interact with F-actin in ABD1a (E171A) or ABD1b (K185A) alone or in both conformations (R221A) (Extended Data Fig. 7a-h). While the K185A mutation (targeting ABD1b alone) had no significant effect, E171A (targeting ABD1a alone) and to a greater extent R221A (targeting both conformations) significantly reduced filopodial density when expressed compared to wild-type (Extended Data Fig. 7d-i).

### ABD1 and ABD2 both interact with multiple F-actin subdomains

The binding footprints of both conformations of ABD1 and ABD2 each not only span two F-actin protomers but also span across multiple actin SDs. The C-terminal regions of both ABD1a and ABD1b contact both SD3 and SD4 of actin protomer A_1_, however ABD1b additionally contacts SD2 (Fig. 2a and b). ABD1a’s N-terminus contacts all four subdomains of actin protomer A_-1_, whilst ABD1b only contacts SD1 and SD3. ABD2 contacts SD1, SD2 and SD3 of protomer A_-1_, while also contacting the D-loop of SD2 in the protomer A_1_ (Fig. 2c).

**Figure 2.**
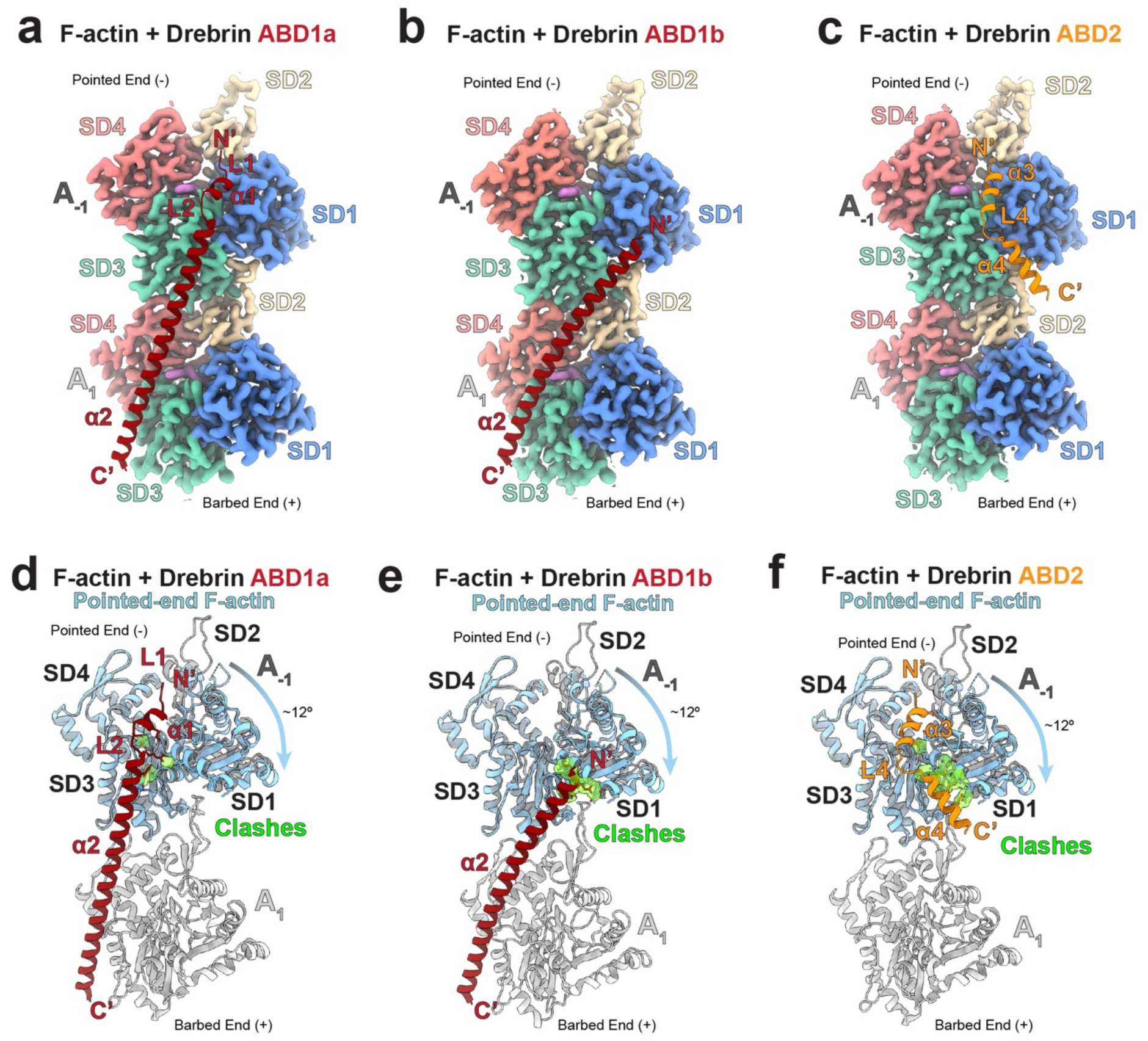
Drebrin’s ABDs bind across multiple actin subdomains. Cryo-EM density for the drebrin ABD-bound F-actin protomer pair (A_-1_ and A_1_) is coloured according to actin subdomain (SD1-4), with the a) ABD1a, b) ABD1b and c) ABD2 models shown to indicate the interaction surface. Superimposition of the ‘twisted’ conformation ultimate pointed end F-actin protomer (light blue, PDB: 9FJO^69^) via SD3 & SD4 onto SD3 & SD4 of actin protomer A_-1_ in our d) ABD1a, e) ABD1b and f) ABD2 models. The ∼12° rotation of SD1 & SD2 relative to SD3 & SD4 in the ‘twisted’ conformation is indicated and resultant clashing regions of this conformation (detected with default settings using ChimeraX’s ‘clashes’ tool^103^) with drebrin ABDs displayed as transparent green density.

Relative to F-actin, G-actin has a ‘twisted’ conformation where SD1 and SD2 are rotated relative to SD3 and SD4^66,67^. This ‘twisted’ conformation also exists at F-actin’s ultimate pointed end protomers, priming them for dissociation^68,69^. Superimposing the cryo-EM structure of an ultimate pointed end F-actin protomer (PDB: 9FJO^69^) via SD3 and 4 onto actin protomer A_-1_ of our drebrin ABD1 or ABD2-bound F-actin structures demonstrates that drebrin’s F-actin binding surfaces are sterically incompatible with the ‘twisted’ conformation (Fig 2d-f).

### Drebrin largely retains its structural features upon F-acting binding

We experimentally assessed the per-residue order/disorder propensity of solution drebrin-E^135–355^ using heteronuclear single quantum correlation (^1^H-^15^N HSQC) nuclear magnetic resonance (NMR) and Chemical shift Z-score (cheZOD)^70^. The amide resonances of drebrin-E^135–355^ ^15^N were assigned using previous NMR studies of drebrin fragments spanning residues 173–238 and 233–317^71,72^ giving 94% coverage of the sequence spanning 173-317 (Extended Data Fig. 8a). The NMR data and the CD analysis (Extended Data Fig. 1a) were broadly in agreement. Under our experimental conditions, cheZOD Z-scores correlated well with secondary structure regions in the F-actin bound state observed by cryo-EM (Extended Data Fig. 8b). This indicates that: 1) F-actin binding does not induce major structural rearrangements to the α-helical organisation in residues 173–317 of drebrin-E^135–355^; and 2) the portions of drebrin’s L3 not resolved in our cryo-EM reconstructions likely remain intrinsically disordered in both apo and F-actin-bound states.

### Overlapping ABD1 and ABD2 binding sites and loop 3 length determine multimodal binding patterns along F-actin

We next investigated how ABD1 and ABD2 may be positioned relative to each other on F-actin. Apart from failing to predict ABD1b altogether, a key error in AlphaFold3’s structural prediction for the drebrin-E^135–355^-F-actin complex was ABD1 helix-α2 being predicted as 26 amino acids longer C-terminally than observed by cryo-EM, such that ABD1a interfaced with three rather than the observed two actin protomers (Extended Data Fig. 1f). Interestingly, secondary structure prediction by JPred4^65^ also predicted a longer helix-α2/shorter L3 than observed (Extended Data Fig. 1c). Instead, experimentally observed length of helix-α2 and L3 by cryo-EM corresponded well with that observed in solution state by NMR (Extended Data Fig. 8b). Therefore, a long L3 (43 amino acids) connecting ABD1 and ABD2 allows them to potentially bind F-actin at several relative positions along the filament simultaneously, including on opposing protofilaments.

In the following analysis we considered ABD1a and ABD1b conformations as one, as their C-terminal E229 from which L3 emanates are roughly found in the same location (<2 Å separation) relative to F-actin (Fig. 1c and e). Several theoretical limitations exist on the permitted relative positions of simultaneously F-actin bound ABD1 and ABD2. Firstly, superimposing our models of ABD1a or ABD1b-decorated F-actin with ABD2-decorated F-actin reveals that neither of the two protomers bound by ABD1 can be co-occupied by ABD2 to avoid steric clashes (Fig. 3a and b). This is consistent with the mutual exclusivity of ABD1 or ABD2 densities on an actin protomer pair seen in our cryo-EM results (Fig. 1b,d and g and Extended Data Fig. 3).

**Figure 3.**
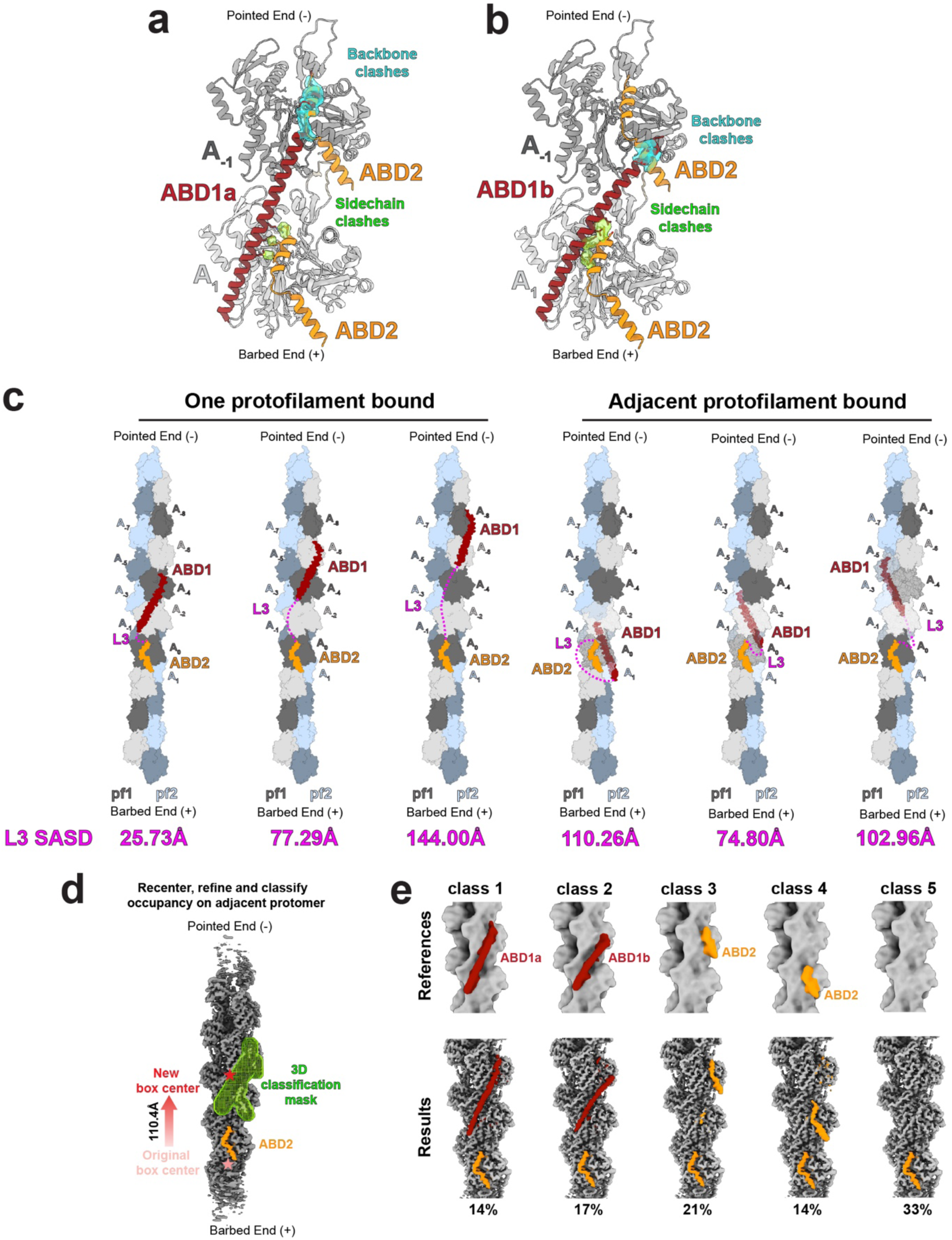
Drebrin’s ABD1 and 2 can adopt a limited number of relative positions along F-actin. Superimposition of the ABD2-bound F-actin model on to each protomer of the a) ABD1a and b) ABD1b-bound F-actin models, showing steric clashes (calculated with ChimeraX’s ‘clashes’ tool^103^ using default settings) with drebrin’s peptide backbone (cyan density) or side chains only (green density). Clashing side chains are displayed. c) Schematic models indicating the 6 permitted positions of ABD1 relative to ABD2 on F-actin. Solvent Accessible Surface Distances (SASDs) determined by JWalk^71^ between the ABD1 C-terminus and ABD2 N-terminus (i.e L3) are shown below each model. d) Schematic showing classification strategy to assess occupancy of ABDs at F-actin substrate sites adjacent to ABD2. Density from the refined reconstruction is shown after re-extracting/recentring/refining 110.4 Å towards the pointed end (new box centre, red) of the original reconstructed ABD2-bound class (original box centre, faded red). The focussed mask used for subsequent 3D classification of ABD occupancy is shown as green mesh. e) Supervised 3D classification references and results, derived from the strategy indicated in panel d. Density is coloured according to molecular identity (F-actin = grey, ABD1 = maroon and ABD2 = orange) and class occupancy % is shown below the results.

Secondly, L3 length cannot exceed a ∼15 nm physical length limit determined by Cα-Cα distance restraints (∼3.5 Å) assuming adoption of an extended linear peptide. However, we must also consider that L3 cannot physically pass through F-actin or other parts of drebrin. We therefore used the program Jwalk^71^ to calculate the shortest distances L3 can cover through solvent across the surface of the protein (solvent accessible surface distance^72^). This resulted in six permitted ABD1 positions relative to ABD2, permitted both sterically and by L3 length, an equal number of which included simultaneous binding the same or adjacent protofilaments (Fig. 3c). Interestingly, in all conformations ABD1 always occupies a protomer pair closer to the F-actin pointed end.

To further investigate whether there is a particular spatial relationship of ABDs on neighbouring protomers along a protofilament, we shifted the centre of our processing 110.4 Å (2x actin protomers) towards the pointed end from a bound ABD2 (Fig. 3d). The data were refined to the new box centre and then supervised classification performed to determine the ABD occupancy on the adjacent protomer. No clear preference was found for ABD occupancies on the adjacent protomer pair (Fig. 3e).

ABD1b’s shifted helix-α2 allows the C-terminus of another ABD1 to bind proximally (closest distance around ∼6 Å, Extended Data Fig.5b and c). As these regions are plentiful in charged/polar residues, we reasoned inter-molecular electrostatics or indirect water-mediated hydrogen bonds could represent a cooperativity mechanism. To investigate, we performed a similar analysis, however instead shifted the centre of processing 55.2 Å (1x actin protomer) towards the pointed end of F-actin relative to a bound ABD1b (Extended Data Fig. 9a). A clear preference for ABD1 binding at this site would suggest ABD1 inter-molecular interactions account for a cooperativity mechanism, however, no such preference was found (Extended Data Fig. 9b). In summary, this analysis suggests drebrin’s two F-actin binding domains can assume up to six relative positions on a single F-actin with no strong preference, due to the long and flexible nature of L3 (Fig. 3c).

### Drebrin is not a strong direct actin-bundling protein

We hypothesised that two drebrin ABDs separated by a flexible L3 could act to bundle F-actin directly. However, the extent of bundling in micrographs of F-actin incubated with drebrin constructs containing either a single or two F-actin binding domains was indistinguishable from F-actin alone (Fig. 4a-e) and in stark contrast to F-actin incubated with fascin-1 (a well characterised bundling protein^73,74^, Fig. 4f). While the distribution of F-actin in micrographs does not preclude a minority of drebrin-E^135–355^ or drebrin-E^1–431^ molecules loosely cross-linking different filaments, the relatively even distribution of filaments across the entire EM grid (data not shown) suggests that drebrin ABD1 and ABD2 predominantly bind the same filament simultaneously.

**Figure 4.**
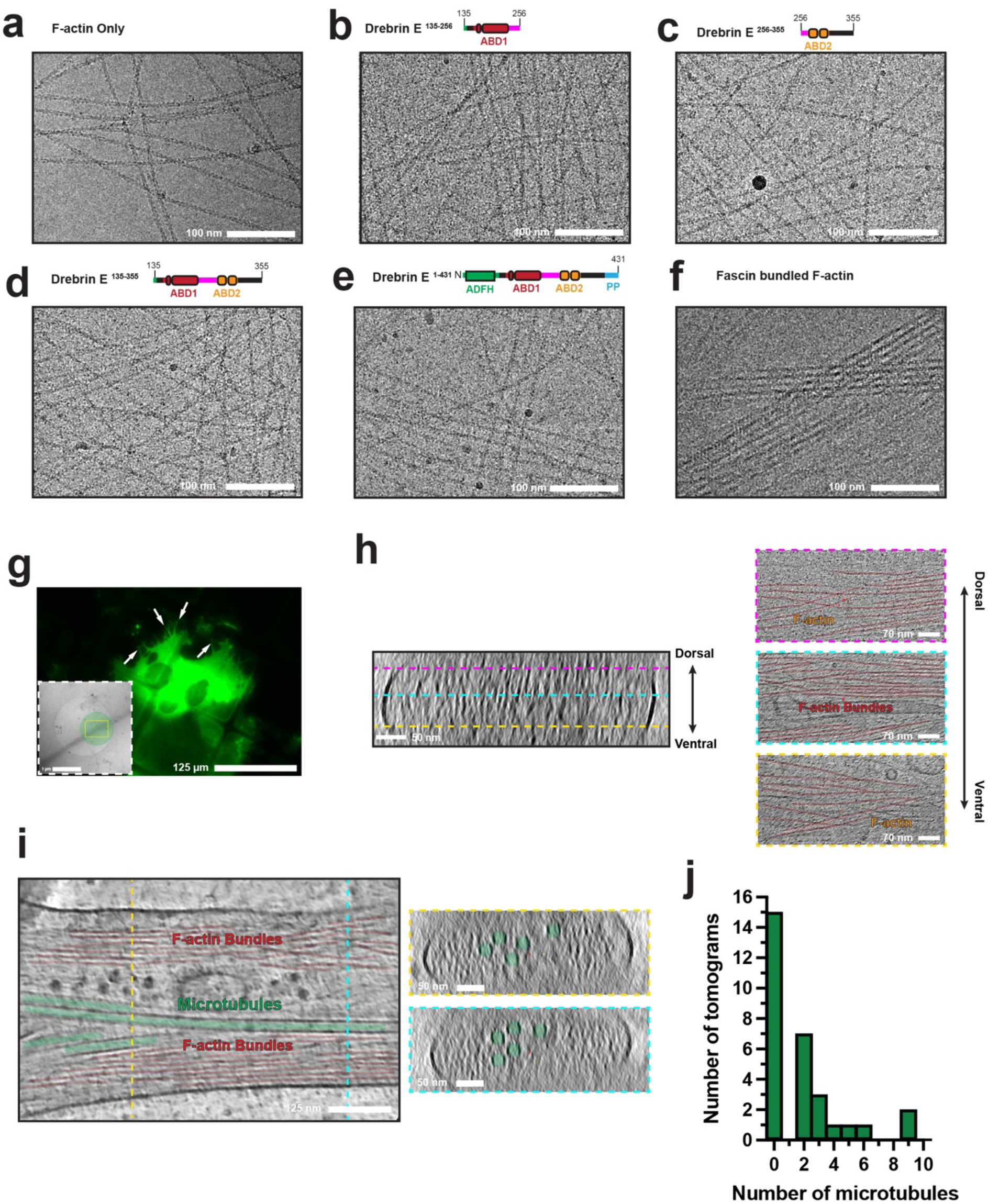
Drebrin is not a direct strong F-actin bundling protein. Micrographs and domain schematics of a) F-actin alone or F-actin decorated with b) Drebrin-E^135–256^, c) Drebrin-E^256–355^, d) Drebrin-E^135–355^, e) Drebrin-E^1–431^ or f) bundling protein fascin-1. g) Light microscopy image of vitrified drebrin-E-eYFP-transfected COS7 cells on EM grids, with arrows indicating drebrin-E-eYFP-overexpressing filopodia. Inset shows a low-mag TEM image of a drebrin-E-eYFP-overexpressing filopodium extending over a hole in the carbon, with the yellow rectangle and green circle indicating the imaged and exposed areas respectively where a tilt series was collected. h) Transverse cryo-electron tomographic section through a drebrin-E-eYFP-overexpressing filopodium (left) with lines indicating the positions of corresponding longitudinal slices through the same tomogram (right) moving from ventral to dorsal positions. i) Longitudinal cryo-electron tomographic section through the medial portion of another drebrin-E-eYFP-overexpressing filopodium (left panel). Transverse sections through the same filopodium, at locations corresponding to the coloured dashed lines in the left panel are shown to the right. In panels h and i microtubules are false coloured in green and an indicative portion of F-actin in red in longitudinal sections. j) Histogram showing the number of microtubules found in 30 tomographic volumes of drebrin-E-eYFP-overexpressing filopodia.

Drebrin’s tendency to induce filopodia when over-expressed in cell lines could hypothetically be derived from a bundling propensity. To test this hypothesis, we transfected COS7 cells (which normally lack filopodia) with drebrin-E-eYFP to induce filopodia as previously described^11^, then collected cryo-tomograms of these filopodia targeted by correlative light and electron microscopy (CLEM), to characterise their underlying F-actin networks (Fig. 4g). F-actin in drebrin-E-induced filopodia was mainly organised into closely packed, collinear bundles, alongside some unbundled or more loosely bundled filaments, particularly in the lateral filopodial periphery (Fig. 4h-i), similar to previous cryo-tomographic observations of filopodia^76,77^. Considering drebrin’s F-actin binding domains are separated by a flexible L3 with a maximum stretched length of ∼15 nm, it seems unlikely drebrin is the core F-actin cross-linker organising these tight bundles. In support of this, constructs containing a single ABD of drebrin induce filopodia comparably to constructs containing both ABDs^12^. Interestingly, in half the drebrin-E-eYFP-transfected filopodia imaged we observed two or more microtubules (Fig. 4i and j), often associated with small organelles, vesicles and ribosomes, which are normally rare features in wild-type filopodia^11,75–77^. This is consistent with previous light microscopy observations showing more common microtubule entry into drebrin-transfected filopodia^11,12^.

### Drebrin modifies the F-actin inter-protomer interface

Actin’s D-loop and C-terminus can adopt various conformations at the inter-protomer interface, associated with F-actin nucleotide state or small molecules such as phalloidin^67,78–80^. In our reconstructions, despite both ABD1 and ABD2 being clearly associating with Mg^2+^-ADP bound F-actin (Extended Data Fig. 10a-c) we observe a closed D-loop and compact C-terminus (Fig. 5a-c) distinct from Mg^2+^-ADP-bound F-actin alone (PDB: 8a2t^67^, Fig. 5d-f and Extended Data Fig. 10d-f). This includes repositioning of key side chains to form inter-protomer interactions, including Q41 on the D-loop and F375 on the extended C-terminus (Fig. 5d-f). This conformation is instead indistinguishable from that observed in the Mg^2+^-ADP-Pi-bound state of F-actin (PDB: 8a2s^67^, Extended Data Fig. 10g-i). Interestingly, only ABD1b and ABD2 directly interact with a portion of the D-loop, while ABD1a does not due to its altered trajectory (Fig. 5a-c). Coincidently, although it takes a similar path, density for the D-loop is poorer in this region and the atomic b-factors are higher in ABD1a-bound F-actin, suggesting these direct interactions are somewhat stabilising (Fig. 5a-c and Extended Data Fig. 10h-j).

**Figure 5.**
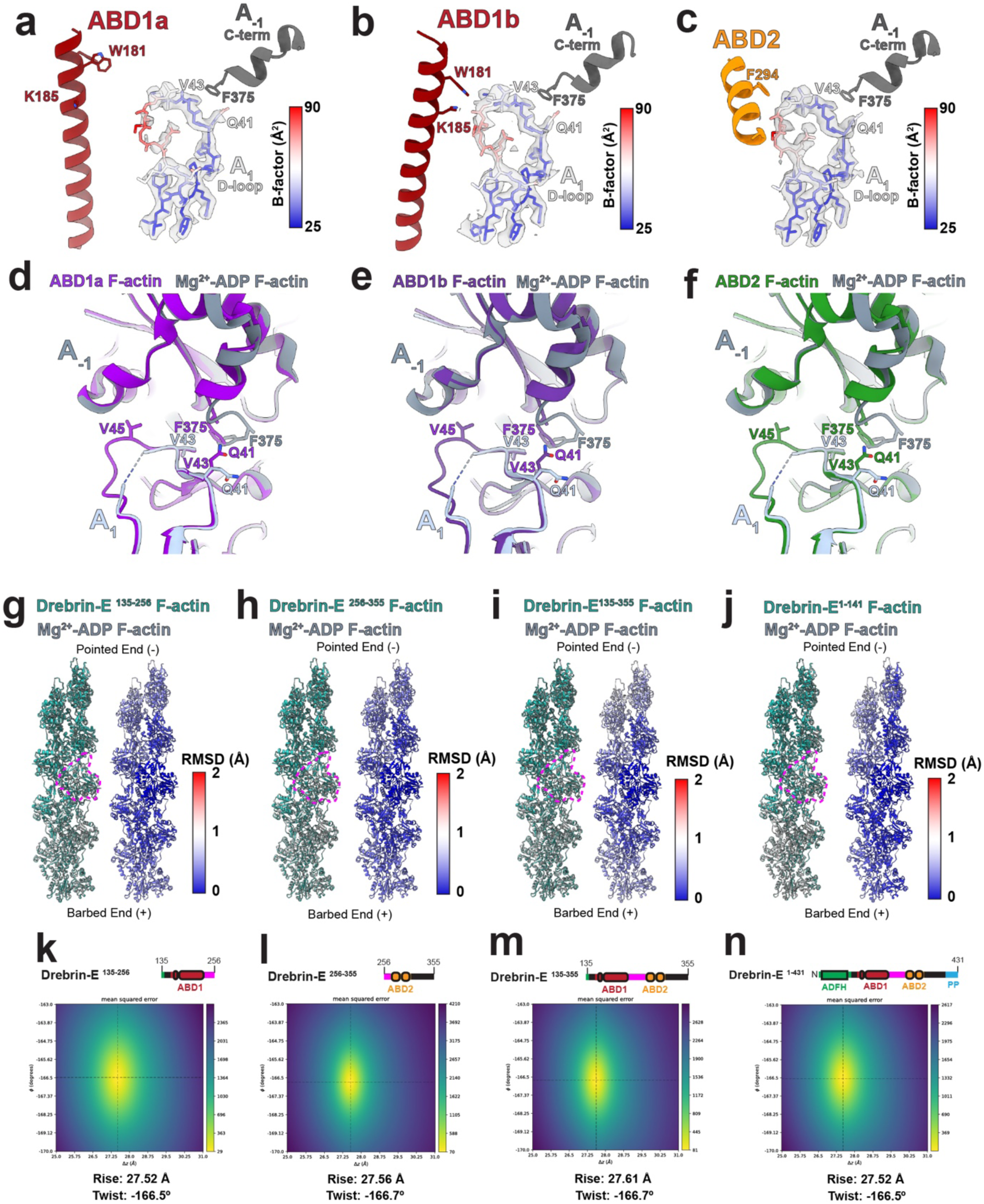
Drebrin modifies the F-actin inter-protomer interface but does not modify F-actin helical rise and twist. Models for a) ABD1a b) ABD1b and c) ABD2-bound F-actin, showing only the ABD and D-loop and C-terminus in proximal actin protomers. The D-loop only is coloured by atomic b-factor. Local-resolution sharpened density for the D-loop is displayed at matching thresholds (RMS level 10) for all three reconstructions. View focussed on the D-loop and C-terminus after superimpositions of our d) ABD1a Mg^2+^-ADP F-actin, e) ABD1b Mg^2+^-ADP F-actin and f) ABD2 Mg^2+^-ADP F-actin models with the Mg^2+^-ADP F-actin alone model (PDB: 8A2T^67^). In all models, key interacting side chains at the inter-protomer and drebrin-D-loop interface are shown (particularly those remodelled in the drebrin-bound state). 9 protomer atomic models were built by fitting Mg^2+^-ADP F-actin protomers (PDB: 8A2T^67^) into drebrin-E^135–256^, drebrin-E^256–355^, drebrin-E^135–355^, drebrin-E^1–431^ or Mg^2+^-ADP F-actin alone (EMD-15106^67^) reconstructions. A central single protomer of these g) drebrin-E^135–256^, h) drebrin-E^256–355^, i) drebrin-E^135–355^ or j) drebrin-E^1–431^ nine protomer F-actin models was then superimposed on a single central protomer of the Mg^2+^-ADP nine protomer F-actin model, to indicate any difference in the helical parameters of repeating protomers along the F-actin axis. The superimposed protomers are indicated with dashed magenta lines. Cα RMSDs are calculated and displayed to the right of the corresponding superimpositions. Helical twist and rise calculated from reconstructions of k) drebrin-E^135–256^, l) drebrin-E^256–355^, m) drebrin-E^135–355^ and n) drebrin-E^1–431^-decorated Mg^2+^-ADP F-actin in CryoSPARC^101^, before any focussed classification of drebrin ABD occupancy. Drebrin-E construct domain schematics are shown above.

### Drebrin does not significantly modify F-actin helical rise and twist or filament curvature

Atomic force microscopy (AFM) studies reported that drebrin increases the cross-over repeat length of F-actin from ∼36 nm to ∼40 nm, proposed to occur via remodelling of F-actin twist^51,52^. We looked for evidence of this remodelling in our cryo-EM data. We first compared 9-protomer F-actin models fitted into drebrin-E^135–256^, drebrin-E^256–355^, drebrin-E^135–355^ and drebrin-E^1–431^-decorated Mg^2+^-ADP F-actin reconstructions prior to focussed ABD classification (Fig. 5g-j) with Mg^2+^-ADP F-actin alone (PDB: 8a2t^67^). When the central protomers were superimposed, we found negligible structural deviation (<0.25 Å) for each protomer rise (helical rise) along the F-actin axis in either direction (Fig. 5g-j), such that the cross-over repeat length would not change more than 3.25 Å.

As an alternative measurement, we then calculated helical parameters (rise and twist) directly from our cryo-EM reconstructions of drebrin-E^135–256^, drebrin-E^256–355^, drebrin-E^135–355^ and drebrin-E^1–431^-decorated Mg^2+^-ADP F-actin before classification of drebrin ABD occupancy and again the differences between these datasets, or that previously reported for Mg^2+^-ADP F-actin alone were negligible (Fig. 5k-n, Mg^2+^-ADP F-actin^67^ = 27.50 Å and 166.7°).

### Mechanisms of drebrin-mediated ABP displacement from F-actin

Our structures allowed us to describe how drebrin displaces ABPs α-actinin, fascin, tropomyosin, myosin and cofilin from F-actin^55,59–62^. Superimposition of F-actin associated with these ABPs onto F-actin in our drebrin ABD1 and ABD2 models revealed clashes that would lead to steric competition.

α-actinin’s actin-binding CH1 domain (PDB: 3LUE^81^) clashes with ABD2 and the ABD1b but not ABD1a conformation (Fig. 6a). Both conformations of drebrin ABD1 and ABD2 share much of their F-actin binding surface with myosin motor domains, such that there are extensive clashes (Fig. 6b). Some of these clashes, such as those with the myosin motor domain HCM-loop, are somewhat dependent on the myosin family member. Fascin has two binding modes with F-actin, allowing F-actin cross-linking^82^. For binding mode 1 (PDB: 8VO5^82^), drebrin’s ABD1b and ABD2 clash with fascin (Fig. 6c), but ABD1a does not. For fascin binding mode 2 (PDB: 8VO6^82^), both conformations of drebrin ABD1 and ABD2 clash with fascin (Fig. 6d). Tropomyosin’s coiled-coil (PDB: 5JLF^83^) has an overlapping binding groove with helix-α2 of drebrin ABD1 (ABD1a more extensively) and therefore abundant clashes are evident (Fig. 6e). ABD2 most likely has minor side chain clashes with tropomyosin, however this is unclear as the only available tropomyosin model lacks side chains. The F-actin-binding ADFH domain of cofilin clashes with both α-helices of ABD2 and both ABD1 conformations (Fig. 6f). Furthermore, our structural data suggests drebrin may additionally inhibit cofilin F-actin interactions through allosteric mechanisms. First, we demonstrate that drebrin’s ABDs stabilise a conformation of F-actin’s D-loop (Fig. 5a-c), which may inhibit cofilin binding^78^. Second, cofilin modifies F-actin from its standard helical structure, inducing a new helical twist and cross-over repeat length via rotations of F-actin subdomains SD1 and SD2^84^. Tight cofilin binding depends on this modification to F-actin^85^. Therefore, by binding over multiple F-actin subdomains and protomers (Fig. 1 and Fig. 2), drebrin’s F-actin binding domains may also act to inhibit F-actin conformational changes associated with cofilin association.

**Figure 6.**
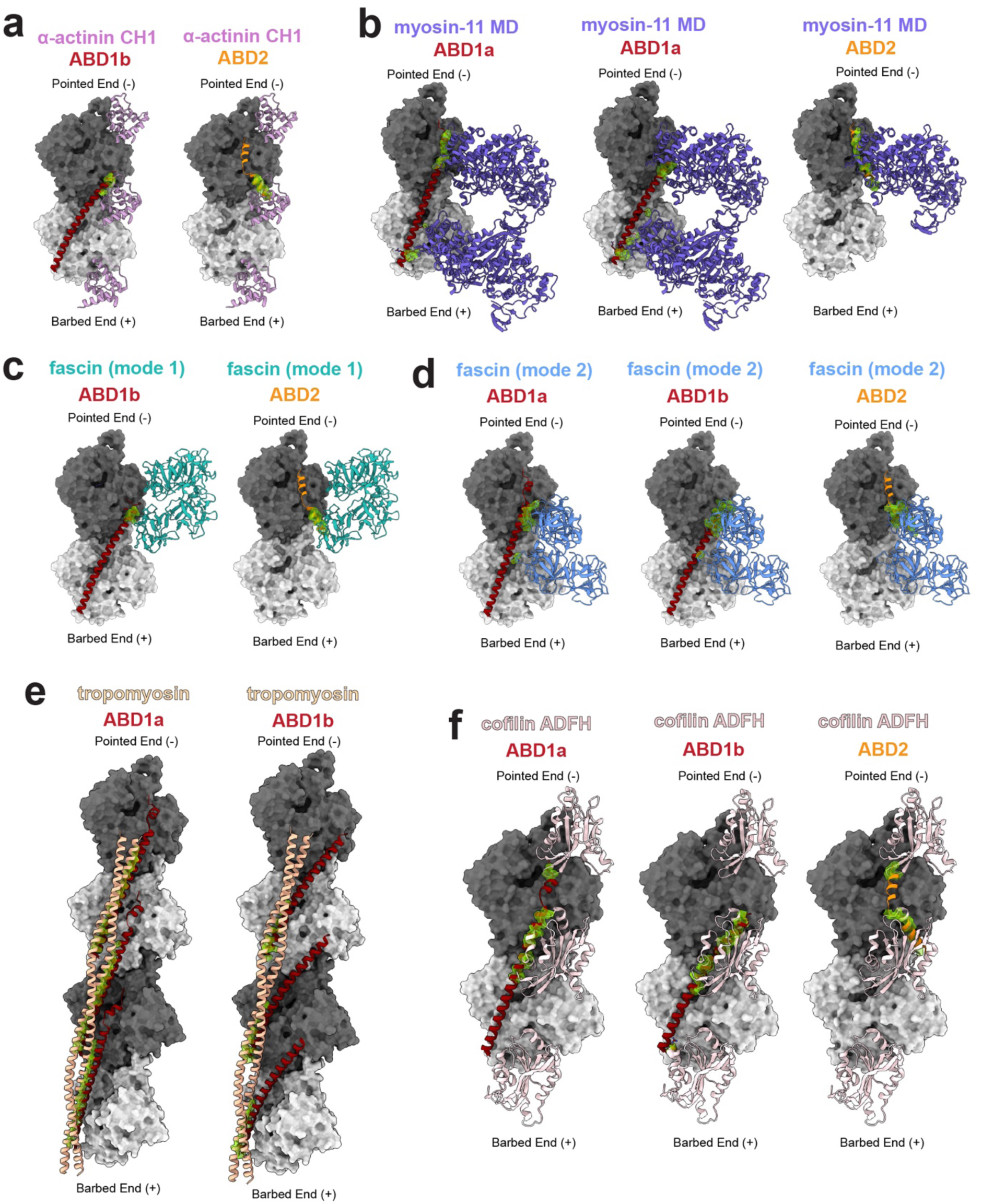
Drebrin sterically competes with key ABPs for F-actin association. Models of F-actin-bound ABPs are superimposed onto our drebrin ABD-bound F-actin models (actin portion as the reference), and resultant clashes (detected with default settings in ChimeraX^103^) shown as green density. Superimposed ABPs; a) α-actinin CH1 domain (PDB: 3LUE^81^), b) myosin-11 motor domain (MD, PDB: 6BIH^115^), c) fascin binding mode 1 (PDB: 8VO5^82^), d) fascin binding mode 2 (PDB: 8VO6^82^), e) tropomyosin (PDB: 5JLF^83^) and f) cofilin ADFH domain (PDB: 3J0S^84^).

## Discussion

In this study we structurally characterise how drebrin interacts with F-actin via two separate ABDs. A re-interpretation of previous affinity binding data of drebrin constructs^2^ suggests that two ABDs confer a 100-1,000-fold increase in affinity for F-actin than one (ABD2) alone. Our structures present no evidence drebrin significantly alters F-actin helical parameters, in contrast with previous AFM measurements^51,52^. With the ABDs competing for an overlapping binding surface, drebrin binds to actin with a stoichiometry of around 1:3, consistent with the later of two reported biochemical assays^2,55^. Because both ABDs bind axially across two protomers and actin subdomain pairs SD1/2 and SD3/4, which are rotated relative to each other in the ‘twisted’ G-actin conformation^66,67^, drebrin can only associate with the F-actin polymer. This ‘twisted’ conformation is also observed at pointed end F-actin protomers favouring their dissociation^68,69^. Therefore, drebrin may stiffen F-actin and prevent its depolymerisation^51–53^ by both cross-linking protomers together and stabilising ‘flattened’ F-actin at pointed ends. This observed binding site contrasts with a previous nanometer resolution negative stain electron microscopy (EM) and cross-linking mass-spectrometry (XL-MS) study which reported globular drebrin densities and major and minor binding sites at different locations on F-actin’s SD1 and SD2^2^. Interestingly, despite the presence of Mg^2+^-ADP in the nucleotide pocket, F-actin’s D-loop and C-terminus were shown to adopt conformations associated with more stable ‘young’ F-actin prior to P_i_ release^67,78^. These conformations were seen at longitudinal protomer interfaces both where drebrin does or does not contact the D-loop (with higher atomic b-factors in non-contacting sites), suggesting drebrin’s effect on these elements could be transmitted by long-range allosteric communication.

Drebrin has a ‘leaky’ F-actin barbed end capping property, which is stronger for the drebrin-A isoform^58,86^. Our structures show the C-terminus of both ABDs, from which flexible sequences emerge, would be located proximal to the barbed end’s apical exposed surface. The drebrin-A-specific insert that has been reported to interact with this barbed end surface^86^ is located after roughly 20 amino acids of flexible sequence following ABD2. Drebrin also interacts with F-actin-nucleating/elongating formins and profilin at barbed ends to modulate F-actin dynamics^58,87,88^. Future studies should aim to further explore the structural bases of drebrin’s modulatory properties at both F-actin ends.

Another way drebrin can modify F-actin dynamics and network architectures in cells is through competing for occupancy with other ABPs. For example, competition of drebrin with cofilin^89^ may augment drebrin’s F-actin direct stabilising properties discussed above in a cellular context. Our structures demonstrate that drebrin’s ability to displace various ABPs from F-actin^55,59–62^ occurs chiefly via steric competition. For all ABPs analysed, both drebrin ABDs to some extent obscure their F-actin binding sites, increasing its competitive potency.

We demonstrate that drebrin’s two ABDs are separated by L3, a loop of 43 amino acids length which is disordered both in solution and when drebrin is bound to F-actin. We therefore anticipated drebrin may cross-link F-actin into bundles, however, there is no clear evidence for strong bundling behaviour in our raw cryo-EM data, consistent with the majority of the literature^55–58^. While the kinetics of drebrin association may favour both ABDs binding the same filament by default, drebrin may participate in F-actin cross-linking in particular instances. For example, by combining drebrin’s F-actin binding function with homer’s tetramerisation, drebrin-homer complexes can mediate F-actin bundle formation^56^. Alternatively, drebrin may participate in cross-linking pre-existing bundles of particular architectures.

Inter-ABD competition and flexible L3 length allows the two ABDs of a drebrin molecule to associate with F-actin via adopting six relative positions along the same or adjacent protofilaments. Interestingly, high sequence conservation and AlphaMissense^90^ pathogenicity scores are observed in the N-terminal portion of L3 despite our cryo-EM maps and solution state NMR showing it as disordered. Recent solution state NMR studies of drebrin constructs containing either ABD1 or ABD2 including the N-terminal portion of L3, also reported this region as disordered^91,92^. This suggests L3’s N-terminus could serve as a protein-binding or post-translational modification site and/or be involved in modifying the F-actin association of ABD1 or its position relative to ABD2. Interestingly, neurofibromin-2 was recently reported to bind to this conserved region of L3, facilitating liver tumorigenesis via the Hippo signalling pathway^46^.

The two ABD1 conformations we observed permit different drebrin stoichiometries on F-actin and, due to their different F-actin interfaces, have different propensities to sterically compete with various ABPs. Mapping sequence conservation onto our models suggests both conformations may play conserved physiological roles, yet their relative prevalence and functional importance in particular contexts are unclear. In the context of filopodial induction after ABD1 overexpression, our mutational analysis suggested ABD1a is more functionally important than ABD1b. However, a mutation targeting both conformations had the strongest effect, which may suggest some adaptive ability for the ABD1b conformation to compensate for mutations targeting ABD1a.

Drebrin (or its F-actin binding region alone) binds F-actin cooperatively, accumulating on F-actin in concentrated clusters^51,52,54^. An inter-molecular cooperativity mechanism could not be determined from our ABD-F-actin structures. Flexible regions of drebrin not observed in our density maps could therefore interact to mediate cooperativity. Alternatively, cooperativity could be mediated through modification to F-actin structure. While drebrin had no effect on F-actin twist or rise, F-actin’s D-loop (and C-terminus) conformation is transmitted along F-actin protomers to unbound sites. Given drebrin ABDs interact directly with this loop, transmission of D-loop conformation could contribute to a cooperativity mechanism.

By utilising two ABDs, one of which can adopt two conformations, and that together can assume variable relative positions on F-actin due to a long flexible connecting loop, drebrin can act as a multi-modal F-actin binder. Furthermore, even within an F-actin protomer pair bound by an ABD, binding of a particular ABP can be sterically permitted only to one of the two protomers. Therefore, this could allow drebrin to accommodate the binding of specific ABPs at sterically permitted sites and even coordinate with other ABPs to form higher order F-actin-based macromolecular architectures, somewhat analogous to the ‘scaffolding’ function of tropomyosin in the sarcomeric thin filament^93,94^. Interestingly, like drebrin, tropomyosin is another example of an ABP that can assume multiple nucleotide-independent positions/orientations of its long α-helical secondary structure upon F-actin, a feature allowing it to adaptively incorporate other ABPs in multimeric complexes with F-actin^95^.

Future work should seek to understand how drebrin’s structure and interaction with F-actin and other binding partners is regulated. For example, there is some evidence phosphorylation of S142, located between the ADFH domain and ABD1 regulates the full-length drebrin fold to modulate ABD1 availability and F-actin cross-linking, as well as exposure of the EB3 binding site^12^. In addition to S142, several other phosphosites including S274 have recently been confirmed as responsive to chemically induced long-term depression^96^. S274 is located at the N-terminal end of ABD2 such that its phosphorylation could modify ABD2 structure or F-actin binding. Interestingly, binding of drebrin to BTP2, an inhibitor which reduces the F-actin-drebrin interaction, is eliminated by the mutation of two proximal lysines just upstream of ABD2; K270 and K271^97^.

In summary, here we have structurally detailed the multimodal interaction of drebrin with F-actin and identified mechanisms of drebrin-mediated F-actin-stabilisation and ABP displacement (Fig. 7a-c). Furthermore, by associating with F-actin via a minor portion of the sequence, drebrin can also serve as platform linking F-actin to structural and signalling proteins, particularly through its remaining flexible or flexibly tethered regions (Fig. 7d). The variability in binding modes may add a complexity to drebrin’s regulation of F-actin architecture and the association of other ABPs and oa-actin binding partners, a subject that warrants future investigation.

**Figure 7.**
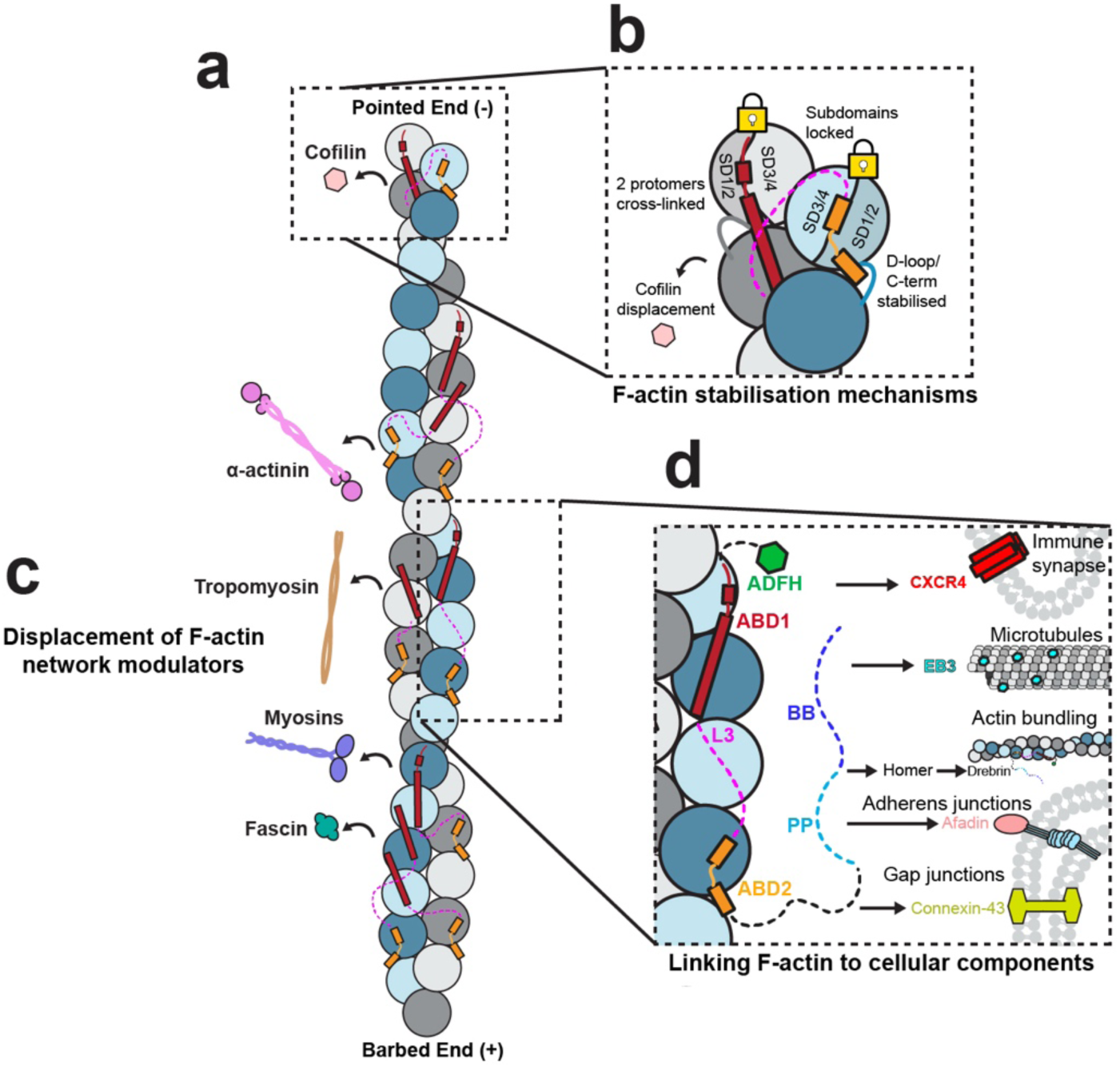
Models of multimodal drebrin-F-actin interactions, F-actin stabilisation and ABP displacement. a) Drebrin ABD1 and ABD2, separated by L3, can assume multiple relative positions on F-actin on a single or adjacent protofilaments. b) Drebrin ABD binding across two actin protomers and their subdomain pairs SD1 & SD2 and SD3 & SD4 cross-links F-actin and stabilises the ‘flattened’ conformation in the ultimate pointed end protomers respectively, preventing depolymerisation. c) ABD association sterically displaces a number of other important ABPs, including cofilin, an actin-depolymerising factor, further contributing to F-actin stabilisation. d) By associating with F-actin via a minor portion of the sequence, drebrin’s ADFH domain, L3, PP domain and BB domain are free to link F-actin to other cellular and molecular components.

## Methods

### Plasmids and site-directed mutagenesis

His-tagged drebrin-E^256–355^ and drebrin-E^135–355^ constructs for protein expression and purification in *Escherichia coli* were previously described and correspond to regions previously characterised as the ‘Hel’ domain or ‘CC and Hel’ domains respectively^12^. Briefly, for drebrin-E^256–355^, human drebrin-E residues 256-355 were subcloned into the pET-24a(+) vector (Novagen). For drebrin-E^135–355^, human drebrin-E residues 135-355 were subcloned into the pET-30a(+) vector (Novagen). For drebrin-E^1–431^ (inclusive of the ADFH domain, regions previously referred to as the ‘CC’ and ‘Hel’ domains and the PP domain^12^), human drebrin-E residues 1-431 were subcloned into the pET-24a(+) vector (Novagen). To make drebrin-E^135–256^ (corresponding to a region previously referred to as the ‘CC’ domain^12^), a stop codon was added to the drebrin-E^135–355^ after position 256 by site-directed mutagenesis using the manufacturer’s instructions alongside manufacturer provided primer designer software (Q5® Site-Directed Mutagenesis Kit, NEB).

The drebrin-E-eYFP (full-length) construct expressed in COS7 cells for CLEM and cryo-ET analysis has been previously described^11^. The drebrin-E^135–256^-eYFP construct used for filopodial induction assays in COS7 cells and as a backbone for site-directed mutagenesis has been previously described^12^. Briefly, for drebrin-E-eYFP and drebrin-E^135–256^-eYFP, human full-length drebrin-E or residues 135-256 respectively were subcloned into the pEGFP-N1 vector (Clontech). Site-directed mutagenesis to produce drebrin-E^135–256^-eYFP mutants for expression in COS7 cells was performed using the drebrin-E^135–256^-eYFP construct as a backbone and the Q5® Site-Directed Mutagenesis Kit (NEB), using the manufacturer’s instructions alongside manufacturer provided primer designer software.

### Protein expression and purification

All drebrin constructs for cryo-EM were expressed in *Escherichia coli* Rosetta (DE3) in ZYM-5052 auto-induction medium^98^. Fascin-1 was expressed in *Escherichia coli* BL21 (DE3) in ZYM-5052 auto-induction medium. Cultures were grown at 37 °C with shaking at 180 rpm to an initial optical density 600 nm (OD₆₀₀) of 0.1, incubated for a further 4 hours, then the temperature decreased to 18 °C for overnight protein expression.

Drebrin-E^135–355^ for NMR studies was ^15^N-labelled by culturing *Escherichia coli* Rosetta (DE3) in M9 minimal media (50 mM Na_2_HPO_4_, 25 mM KH_2_PO_4_, 10 mM NaCl, 20 mM ^15^NH_4_Cl, 20 mM D-glucose, 0.3 mM CaCl_2_, 1 mM MgSO_4_, 1 mg thiamine hydrochloride, 1 mg biotin). Cells were grown at 37 °C to an optical density 600 nm (OD₆₀₀) of 0.9 then induced with 1 mM IPTG with shaking at 18 °C overnight for protein expression.

Cell pellets were lysed by sonication with buffer containing 20 mM HEPES, pH 7.5, 500 mM NaCl, 10% (w/v) glycerol, and EDTA-free protease inhibitor. For all constructs, lysates were clarified by centrifugation at 48,400 x g for 30 minutes at 4 °C and loaded onto a HiTrap TALON® crude column (Cytiva) for purification by immobilised metal-ion affinity chromatography (IMAC). The column was washed with IMAC wash buffer (20mM HEPES, pH 7.5, 500 mM NaCl). Bound protein was eluted with IMAC elution buffer (20mM HEPES, pH 7.5, 500 mM NaCl, 250 mM imidazole), then buffer exchanged into ion-exchange (IEX) binding buffer (20mM HEPES, pH 7.5, 50 mM NaCl, 2 mM DTT) using a HiPrep 26/10 Desalting column (Cytiva). Protein in IEX binding buffer was then loaded onto a HiTrap Q HP anion exchange chromatography column (Cytiva), then eluted in a continuous gradient from IEX binding buffer to IEX elution buffer (20mM HEPES, pH 7.5, 1 M NaCl, 2 mM DTT). Finally, protein was further purified by size exclusion chromatography using a Superdex 200 Increase 10/300 GL (Cytiva) in HEPES-KME buffer (10 mM HEPES, pH 7.5, 50 mM KCl, 1 mM MgCl_2_, 1 mM EGTA, 1 mM DTT).

### Sample preparation for single-particle cryo-EM

Actin was reconstituted from lyophilised rabbit skeletal muscle actin powder (Cytoskeleton, Inc.) on ice to 10 mg/ml (5 mM Tris-HCl pH 8.0, 0.2 mM CaCl_2_, 0.2 mM ATP, 5% (w/v) sucrose, and 1% (w/v) dextran), then snap frozen with liquid nitrogen and stored at –80 °C. Reconstituted G-actin was centrifuged at 12,000 x g at 4 °C to remove aggregates. To polymerise F-actin, the supernatant was diluted to 1 mg/ml actin with HEPES-KME buffer with the addition of 0.2 mM ATP. Polymerised F-actin was stored at 4 °C for at least 24 hours before use.

Immediately before use, F-actin was diluted in HEPES-KME buffer to a final concentration of 8 µM. Drebrin constructs were diluted to 72 µM and fascin-1 diluted to 15uM in the same buffer. 3 µl of F-actin was applied to glow-discharged EM grids and incubated for 1 min at room temperature. Excess buffer was manually blotted away, followed by the addition of 3 µl drebrin or fascin-1 solution and incubation for 1 min at room temperature. Excess solution was again blotted away, and a second 3 µl of drebrin or fascin-1 solution was applied. Grids were then transferred to an EM GP2 plunge freezer (Leica Microsystems), incubated for 1 min at 25 °C and 80% relative humidity, front blotted for 6 seconds and plunge-vitrified in liquid ethane. For preparation of F-actin with drebrin-E^135–355^, Au-flat™ gold EM grids with gold alloy film (1.2 µm holes, 1.3 µm spacing, Electron Microscopy Sciences) were used. For preparation of drebrin-E^135–256^, drebrin-E^256–355^, drebrin-E^1–431^ and fascin-1 with F-actin, C-flat™ carbon-coated copper EM grids (1.2 µm holes, 1.3 µm spacing, Protochips) were used.

### Single-particle cryo-EM data collection

Datasets of drebrin-E^1–431^, drebrin-E^135–256^ and drebrin-E^256–355^ with F-actin were collected using a Krios G3i operating at 300 kV, with a Gatan K3 direct electron detector and a Bioquantum Imaging Filter. Low-dose movies were collected automatically with EPU (ThermoFisher) software at 0.825 Å/pixel, in zero-loss imaging mode with a 20 eV energy-selecting slit. A defocus range of 0.7–2.5 μm was used, with a total dose of 64 e−/Å^2^ spread over 59 frames in electron counting mode (5 e-per pixel per second). Datasets of drebrin-E^135–355^ with F-actin were collected using a Krios III operating at 300 kV, with a Falcon 4i direct electron detector and a SelectrisX energy filter. Low-dose movies were collected automatically with EPU (ThermoFisher) software at 0.931 Å/pixel, in zero-loss imaging mode with a 10 eV energy-selecting slit. A defocus range of 0.7–2.5 μm was used and the total movie dose was 45 e−/Å^2^ spread over 44 frames in electron counting mode.

### Cryo-EM image processing

Pre-processing steps were identical for all datasets. Movies were gain-corrected, motion-corrected, dose-weighted, and summed using RELIONv5’s implementation of MotionCor2^99^. CTF determination was then performed on micrographs using CTFFIND4^100^, then micrographs and metadata imported into CryoSPARC^101^ for particle picking and initial refinements.

For all datasets, F-actin filaments were picked with an inter-box distance of 27.6 Å (corresponding to the helical rise of an actin protomer) using CryoSPARC’s filament tracer^101^ in template-free mode. Picked particles were curated based on normalised cross-correlation, power score, filament curvature, and filament sinuosity. For all datasets apart from F-actin with drebrin-E^256–355^ selected particles were extracted and 4x binned at this stage. For the dataset of F-actin with drebrin-E^256–355^ selected particles were extracted unbinned. Particles were then subjected to 2D classification in CryoSPARC^101^ to remove poor particles. Good classes were re-centred, 2x binned for all datasets apart from F-actin with drebrin-E^256–355^ (which was kept unbinned) and used for an initial 3D refinement in CryoSPARC^101^ using the helical refinement job with a helical symmetry order of 1. The 3D refinement reference volume was generated from PDB ID:7BT7^102^ (F-actin-Mg²⁺-ADP) using the molmap command in ChimeraX v1.9^103^ and then low-pass filtered to 20 Å resolution. Processing of the drebrin-E^1–431^ with F-actin dataset was terminated at this stage, whereas all other datasets underwent further processing.

Duplicate particles were removed, and particle coordinates were converted into RELIONv5^99^. star files using PyEM (https://github.com/asarnow/pyem). Unbinned (drebrin-E^256–355^ with F-actin dataset) or 2x binned particles (drebrin-E^135–256^, and drebrin-E^135–355^ with F-actin datasets) were then re-extracted in RELIONv5^99^ and locally refined with C1 symmetry to correct potential orientational and translational offsets introduced during metadata conversion.

The drebrin-E^135–256^ and drebrin-E^256–355^ datasets were first examined to identify conformations of ABD1a, ABD1b, and ABD2. Initial explorative processing with unsupervised 3D classification revealed rough densities representing ABD1a and ABD1b in the drebrin-E^135–256^ dataset, and ABD2 in the drebrin-E^256–355^ dataset. Based on ABD1 densities, masks and reference volumes for ABD1a and ABD1b were generated by trimming away all but ordered regions of ABD1 from the AlphaFold3-predicted models of drebrin-E^135–355^-F-actin, rigid-fitting these into density alongside F-actin protomer models (PDB:7BT7^102^, F-actin-Mg²⁺-ADP) then using the molmap command in ChimeraX^103^ and low-pass filtering the resulting density to 10 Å resolution. Then for the drebrin-E^135–256^ dataset, ABD1a or ABD1b-occupied actin protomer pairs were separated by focussed supervised 3D classification using these references without alignment in RELIONv5^99^ using and a T-value of 250. A soft classification mask was made to include both possible ABD1 conformations on the central actin protomer pair. Resulting ABD1a and ABD1b classes were subsequently each locally refined with C1 symmetry using a soft mask encompassing 35% of the filament.

For the drebrin-E^256–355^ dataset, our initial explorative processing with unsupervised 3D classification revealed ABD2 binds F-actin on every actin protomer (a stoichiometry of 1:1). Therefore, unbinned particles extracted in RELIONv5^99^ were not classified but instead subjected 2 rounds of local refinement each followed by CTF refinement and Bayesian polishing, followed by a final local refinement. Local refinements used a soft mask encompassing 35% of the filament.

A similar supervised classification strategy was applied to the drebrin-E^135–355^ dataset. However, in addition to using the ABD1 reference maps described above for drebrin-E^135–256^ F-actin dataset processing, additional reference maps representing ABD2 bound to the upper, lower, or both actin protomers were created. These references were made from preliminary ABD2 models with actin protomers derived PDB:7BT7^102^ (F-actin-Mg²⁺-ADP) built into the final reconstruction from the drebrin-E^135–256^ dataset, using the molmap command in ChimeraX v1.9^103^ and low-pass filtering to 20 Å resolution. The corresponding reference densities were included within a soft focussed mask (Extended Data Fig. 3). Classes corresponding to ABD1a, ABD1b, or ABD2 bound F-actin in the correct register were selected and unbinned and re-centred particles extracted. Each class was then subjected to an initial local C1 3D refinement, followed by CTF refinement and Bayesian polishing, followed by a subsequent C1 local refinement. All local 3D refinements up to this point used a soft Z mask encompassing 35% of the filament. For the ABD1a and ABD1b classes, an additional focussed local refinement was performed using a soft mask containing two actin protomers and the corresponding ABD1 density. Local refinements in RELIONv5^99^ used an initial angular search range of +/-3° and an initial offset range of +/-3 pixels.

Global resolutions from independently refined half-maps were estimated using the gold-standard Fourier shell correlation (FSC) 0.143 cutoff criterion (noise-substitution test–corrected^104^) and the soft masks applied during the final refinement steps. As an additional resolution measure, FSCs were calculated between maps and models in Phenix (v1.21.2-5419)^105^ and resolutions estimated using the FSC 0.5 cutoff criterion. Local resolution was estimated in RELIONv5^99^. Displayed cryo-EM densities are locally sharpened with DeepEMhancer^106^ or local resolution filtering in RELIONv5^99^ as stated.

### Pseudo-atomic model building

Initial models of drebrin ABDs bound to F-actin were generated using AlphaFold3 by providing two copies of human drebrin-E^135–355^ together with six copies of human α-skeletal muscle actin and six copies each of ATP and Mg²⁺ as ligands. Drebrin ABD1a, ABD1b and ABD2 starting models were then produced by adjusting the position then rigid fitting (using the fit-in-map tool in ChimeraX^103^) regions of the predicted drebrin model into cryo-EM density, then removing parts of the model which lacked corresponding density.

The actin chains in the predicted assembly were replaced by fitting the central two actin protomers from the Mg²⁺-ADP-Pi F-actin structure (PDB 8a2s^67^) into the maps and removing the Pi. Iterative rounds of model adjustment in Coot (v0.9.8.92)^107^ and real-space refinement in Phenix (v1.21.2-5419)^105^ were subsequently performed, using density maps sharpened according to local resolution in RELIONv5^99^.

### Cellular cryo-ET sample preparation

Gold H2 London finder EM grids with overlaid holey carbon (3 μm holes with 3 μm spacing, Quantifoil) were placed on the surface of 35 mm cell culture dishes (Nunc), then sterilised by UV treatment for 20 minutes on each side.

COS7 cells (ECACC 87021302) were plated in 6-well culture plates (NunclonTM Delta Surface, ThermoScientific 140675) and maintained at 37°C in 5% CO_2_ in humidified air in Dulbecco’s Modified Eagle Medium + GlutaMax (Gibco, 31966-021) containing 10% heat-inactivated foetal bovine serum (Gibco BRL, 16140-071) and 100 I.U./ml penicillin/100 I.U./ml streptomycin (Sigma, P0781). Cells at >90% confluency were transfected in Opti-MEM I medium (Gibco, 51985-034) using Lipofectamine 2000 (Invitrogen, 11668027) and 4μg of drebrin-eYFP DNA plasmid^8^ per well. Cells were left for 48 hours at 37°C and 5% CO_2_, released by trypsinisation then re-plated on the pre-prepared EM grids overlaid on 35mm cell culture dishes. After another 24 hours, cells were vitrified using back-sided blotting (8-12 second blot time) on an automated Leica GP plunge freezing device operating at 37°C and 75% humidity.

### Cryo-fluorescence microscopy and cryo-ET

Imaging of cellular drebrin-E-eYFP fluorescence was performed on unclipped grids maintained in the vitreous cryogenic state within a Linkam stage (CMS196) operating on a Nikon Eclipse E200 fluorescence microscope. Images were acquired using a 20x air-objective lens and stitched into a full EM grid map using NIS elements software.

For cryo-ET, grids were then clipped and transferred to a ThermoFisher Titan Krios G3i operating with a Gatan K3 direct detector and BioQuantum energy filter with a 20 eV energy-selecting slit. Cryo-fluorescence grid maps were correlated with low-mag transmission cryo-EM grid maps by manual overlay and target drebrin-E-eYFP transfected COS7 cell filopodia identified. Tilt series were acquired at target sites using a dose-symmetric tilt^108^ scheme from – 60 to +60° with 3° increments and a total dose of 120 electrons/Å^2^. Each tilt movie with 3 fractions was acquired in super-resolution mode using a sampling of 2.1Å/physical pixel with a dose of 2.92 electrons/Å^2^.

Movies of individual tilts were gain-corrected, motion-corrected, dose-weighted, and summed using RELIONv5’s^99^ implementation of MotionCor2^99^. CTF determination was then performed on tilt images using CTFFIND4^100^. Tilt-series were aligned using AreTomo2^109^ wrapper within RELION v5^99^, with an estimated tomogram thickness oa 200 nm, and bad tilts excluded by visual inspection. Tomograms were reconstructed using RELIONv5^99^ with dimensions of 4092ξ5760ξ2000 and binned x5 (to 10.5 Å/pixel final pixel size). Tomograms for display were denoised using IsoNET^110^.

### Circular Dichroism (CD) spectroscopy

CD spectra were acquired at 23°C on Applied Photophysics Chirascan Plus spectrometer (Leatherhead, UK), using 0.5 mm path-length Quartz Suprasil rectangular cells. Data collection parameters were set to 2 nm spectral bandwidth, 1 nm step size, and 1 s instrument time per point. Spectra for drebrin^1–431^, drebrin^135–355^, drebrin^135–355^ ^15^N were collected at concentrations of 0.170, 0.077, 0.107 mg/ml respectively. All spectra were adjusted by light-scattering, buffer subtracted, smoothed, corrected for protein concentration and expressed in terms of mean residue ellipticity (MRE). Secondary structure content was predicted using BeStSel web server^111^.

### Nuclear Magnetic Resonance (NMR)

NMR studies were performed using a Bruker Avance spectrometer operating at a proton frequency of 800 MHz. Heteronuclear single quantum correlation (^1^H-^15^N HSQC) experiments were conducted at 293.15 K using a ^15^N-labelled sample of drebrin-E^135–355^ (^15^N-drebrin-E^135–355^) at a final concentration of 50 μM in 17 mM NaH_2_PO_4_, 3 mM Na_2_HPO_4_, 50 mM NaCl, pH 6. NMR spectra were processed using TopSpin® (Bruker) and analysed with CcpNmr V3^1^^12^. The assignments of the region between K173 and R317 was transferred from the Biological Magnetic Resonance Data Bank entries 52895 and 52729^91,92^. Order/disorder propensity was predicted with cheZOD^70^, using the chemical shift perturbations of the NMR assignment. Data were analysed and displayed using Microsoft Excel and Prism 7 GraphPad (Dotmatics).

### Sequence alignment of drebrin’s actin-binding region

131 human drebrin orthologue canonical vertebrate protein sequences were downloaded from Ensembl release 115^1^^13^. 93 sequences have a 1-to-1 relationship with human drebrin and 38 fish sequences have a 1-to-many relationship with human drebrin. Sequences were iteratively aligned using MUSCLE software, with default parameters^114^, trimmed and curated to Homo sapiens drebrin residues 135-308 (common to human isoforms and containing both ABDs). The final sequence alignment retained drebrin sequences from 114 species including Homo sapiens.

### Cell Culture and Confocal Imaging

COS7 cells (ECACC 87021302) were plated in 24-wells plates (Nunclon^TM^ Delta Surface, ThermoScientific) with coverslips and maintained at 37°C in 5% CO_2_ in humidified air in Dulbecco’s Modified Eagle Medium, no glucose (Gibco, 11966025) containing 10% fetal bovine serum (Gibco, A5256701) and 100 I.U./ml penicillin/100 I.U./ml streptomycin (Sigma, P0781). Media was changed for antibiotic-free media (recipe as above but with antibiotics) before transfection. Cells at >70% confluency were transfected using Lipofectamine 2000 (Invitrogen, 11668027) following the manufacturer’s instructions. A mix of 500ng of DNA plasmid and 2µl of Lipofectamine 2000 reagent diluted in Opti-MEM^TM^ (ThermoScientific) was used to transfect each well. Transfected cells were incubated for 18 hours, then the Lipofectamine removed by a change into fresh media containing antibiotics, followed by another 24-hour incubation. Coverslips with transfected cells were fixed with 4% paraformaldehyde then permeabilised with PBS containing 5% (w/v) bovine serum albumin and 0.2% (v/v) Triton X-100. Coverslips were then stained with phalloidin Alexa Fluor™ 568 (Invitrogen, A12380) and mounted on glass slides using ProLong™ Glass Antifade Mountant (Invitrogen, P36982).

Cells were imaged on a Nikon AXR inverted confocal microscope equipped with a 60× Plan Apo Lambda D oil-immersion objective. Finger-like protrusions extending beyond the cell membrane were identified as filopodia manually, and those <1 µm in length (measured from tip to base) in addition to those located along concave regions of the membrane (retraction fibres) were excluded from final quantification. Cell perimeter distances were measured using FIJI.

## Data availability

Focussed reconstructions of ABD1a, ABD1b and ABD2 bound to an F-actin protomer pair from the drebrin-E^135–355^-decorated F-actin data set are deposited in the Electron Microscopy Data Bank (EMDB) alongside corresponding atomic models deposited in the Protein Data Bank (PDB); F-actin-ABD1a: EMD-55893 and PDB: 9TG8, F-actin-ABD1b: EMD-55894 and PDB: 9TG9 and F-actin-ABD2: EMD-55895 and PDB: 9TGA.

## Contributions

W.Z, G.A, T.M, and N.N carried out the experiments. J.A. and W.Z. conceived the study, designed the experiments, interpreted the results, and authored the paper. F.O and P.R.G-W produced and provided initial wild-type drebrin-E expression constructs and drebrin-E-eYFP transfected COS7 cells. M.R.C, T.M and P.R.G-W interpreted the results and contributed to revisions of the paper.

## Ethics declaration

Competing interests: the authors declare no competing interests.

## Acknowledgements

This work was supported by a China Scholarship Council (CSC) PhD studentship awarded to W.Z, BBSRC research grants awarded to J.A (BB/V006568/1 and UKRI2979) and a Leverhulme Trust grant (ref: RPG-2020264) awarded to M.R.C and G.A. The Topf lab (M.T, T.M and N.N) thank the Leibniz Institute of Virology as part of Leibniz Science Campus InterACt (funded by the BWFGB Hamburg and the Leibniz Association), a Wellcome Collaborative Award in Science (209250/Z/17/ Z) and the Landesforschungsförderung Hamburg (HamburgX). P.R.G-W was supported by a Leverhulme Trust Emeritus Fellowship (ref: EM-2022-038\2).

We thank the London Consortium for Electron Microscopy (LonCEM), which supported by Wellcome Trust grant 206175/Z/17/Z and its partner institutes. We acknowledge Diamond for access and support of the cryo-EM facilities at the UK national electron Bio-Imaging Centre (eBIC), proposal BI38847. We acknowledge the KCL centre for ultrastructural imaging (CUI) for access and support with grid vitrification, cell culture and cryo-fluorescence microscopy. We thank the Nikon Imaging Centre at Kings College London for assistance with confocal microscopy experiments. We acknowledge the NMR and CD facilities at the Centre for Biomolecular Spectroscopy at King’s College London, funded by the Wellcome Trust and British Heart Foundation (ref. 202767/Z/16/Z and IG/16/2/32273 respectively to Maria R Conte and others). We thank Tam Bui for assistance with CD experiments. We thank Dr Michelle Simon at the Hub for Applied Bioinformatics (HAB), King’s College London for assistance with sequence bioinformatics.

## Extended Data Figures and Figure Legends

**Extended Data Fig. 1.**
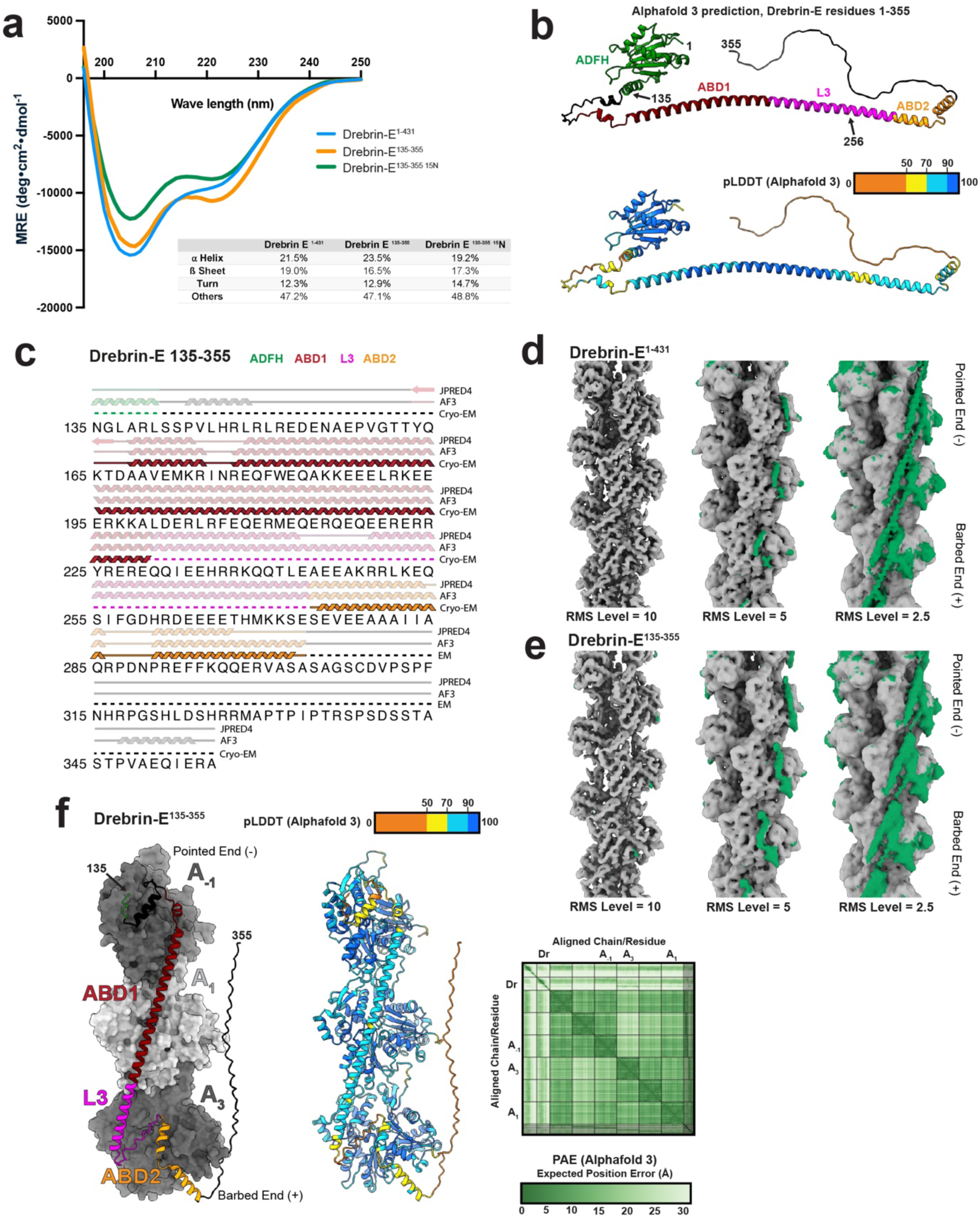
Initial analysis of drebrin-E constructs and F-actin interactions. a) Circular dichroism spectra of drebrin-E^1–431^, drebrin-E^135–355^ and ^15^N-drebrin-E^135–355^ (used in NMR studies). The inset shows a table indicating secondary structure content prediction by the BeStSel server^1^^11^. b) Structural prediction of drebrin-E residues 1-355 with Alphafold3^64^, coloured by domain with amino acids at construct boundaries indicated (top panel) or pLDDT confidence score (bottom panel). c) The sequence for drebrin-E amino acids 135-355 is shown, with secondary structure predicted by JPred4^65^ or Alphafold3^64^ compared with that found by cryo-EM (ABD1a and ABD2) shown above. Predictions are shown as faded (compared to experimental data) and dashed lines indicate regions unresolved due to flexibility in our cryo-EM reconstructions. Initial cryo-EM reconstructions of d) drebrin-E^1–431^ and e) drebrin-E^135–355^, before image classification, shown at three different threshold RMS levels as indicated. The Mg^2+^-ADP bound F-actin model (PDB: 8a2t^67^) was fitted into the density and the colour Zone function in ChimeraX^103^ used to colour density 5 Å from the fitted model green, indicating density not accounted for by actin. f) Structural prediction of two drebrin-E^135–355^ molecules with 6 Mg^2+^-ADP-bound F-actin protomers with Alphafold3^64^ was performed. The highest scoring drebrin-E^135–355^ molecule and its three interacting actin protomers are shown, coloured by domain (left) or pLDDT confidence score (right). The predicted aligned error (PAE) plot for this prediction is shown below, with chains shown above indicated on the X and Y axes (Dr = drebin-E^135–355^, A = actin protomer).

**Extended Data Fig. 2.**
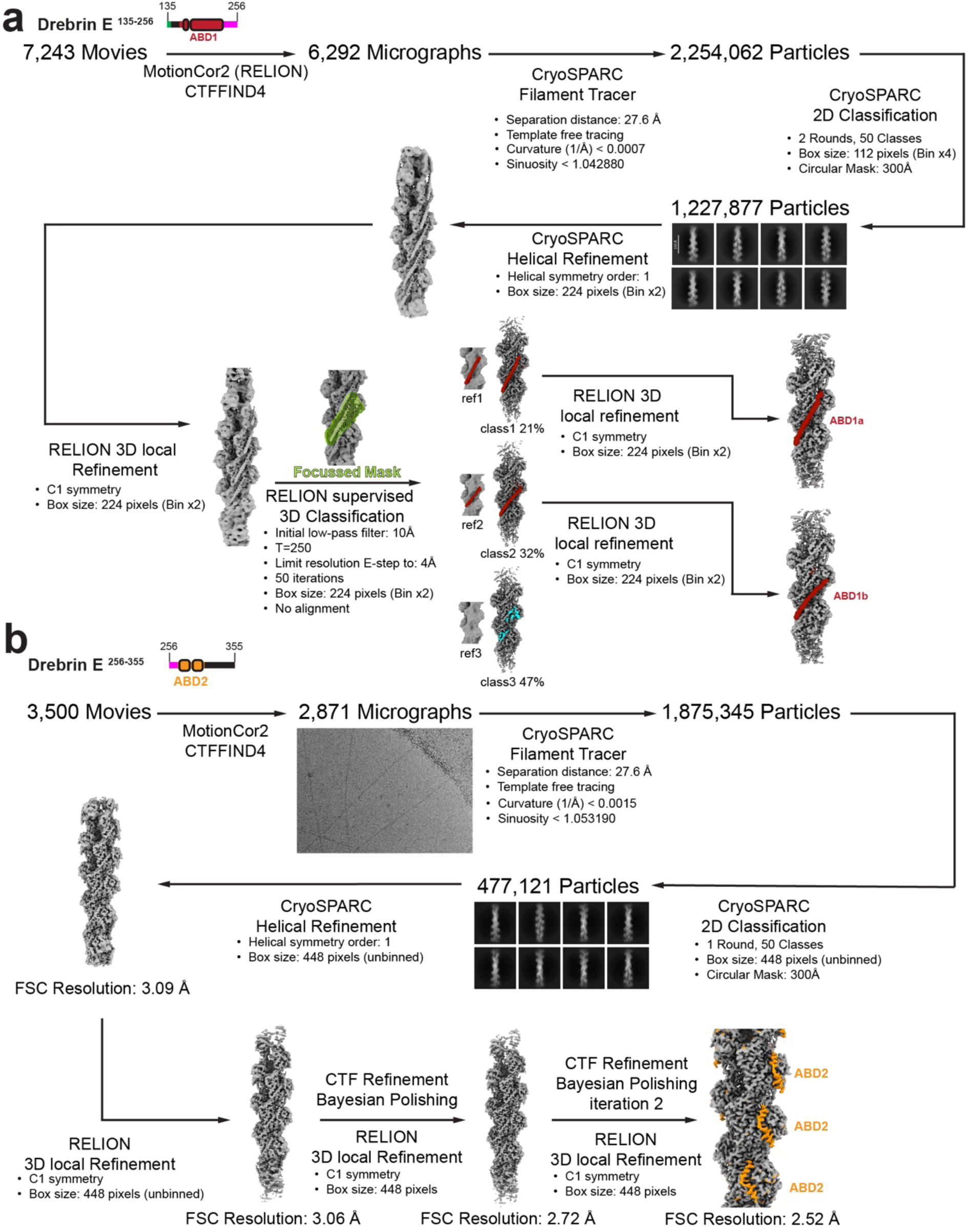
Image processing strategies and ABD1 or ABD2 reconstructions for F-actin decorated with drebrin-E^135–256^ or drebrin-E^256–355^ respectively. a) Schematic indicating the image and particle processing strategy for our dataset of F-actin decorated with a) drebrin-E^135–256^ and b) drebrin-E^256–355^ including initial steps in CryoSparc v4.6.2^101^ and focussed classification and refinement steps in RELIONv5^99^. Cryo-EM reconstructions displayed are unsharpened densities.

**Extended Data Fig. 3.**
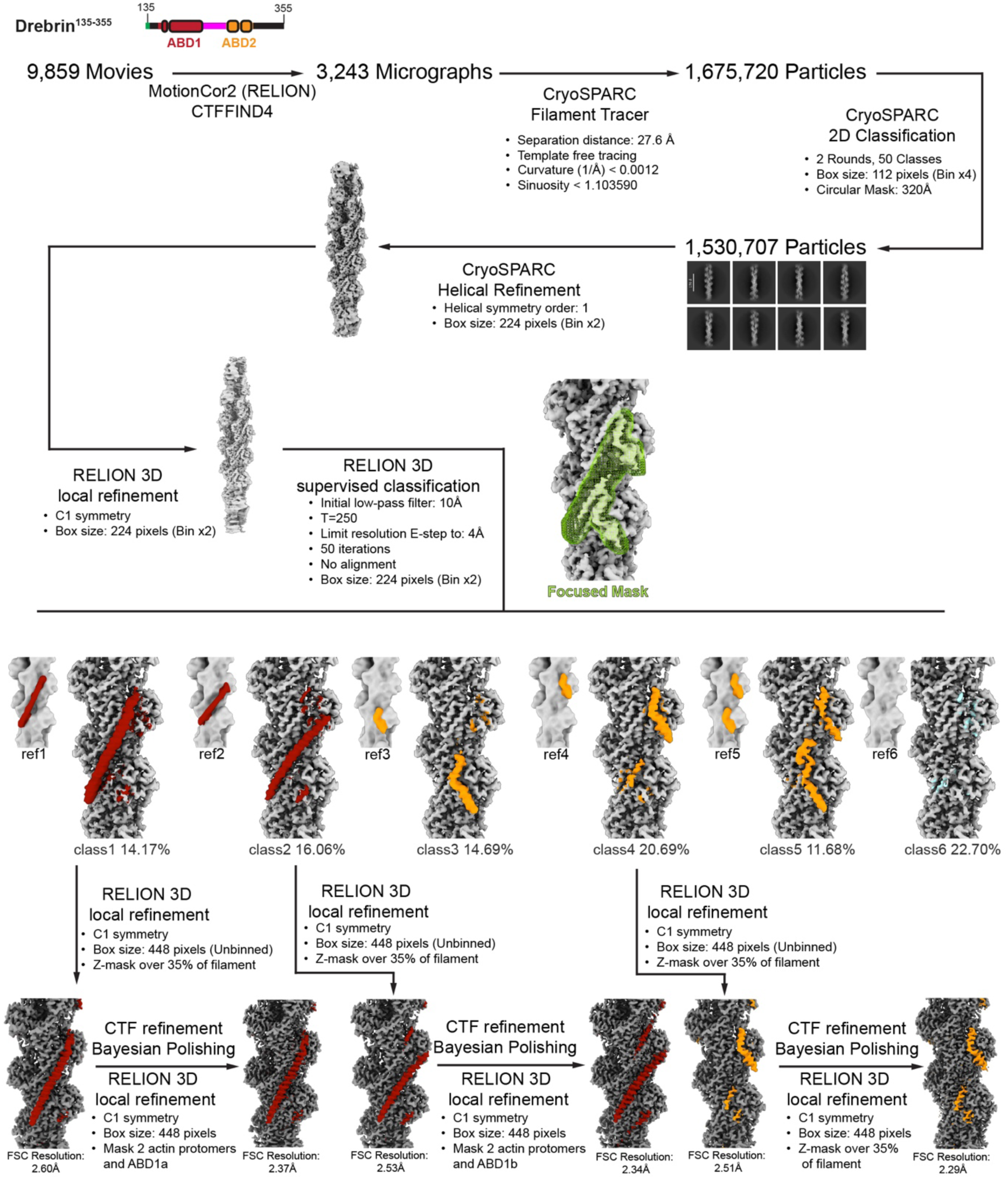
Image processing strategy for F-actin decorated with drebrin-E^135–355^. Schematic indicating the image and particle processing strategy for our dataset of F-actin decorated with drebrin-E^135–355^, including initial steps in CryoSparc v4.6.2^101^ and focussed classification and refinement steps in RELIONv5^99^. Displayed final reconstructions were locally sharpened according to local resolution with RELIONv5^99^, all prior displayed densities are unsharpened.

**Extended Data Fig. 4.**
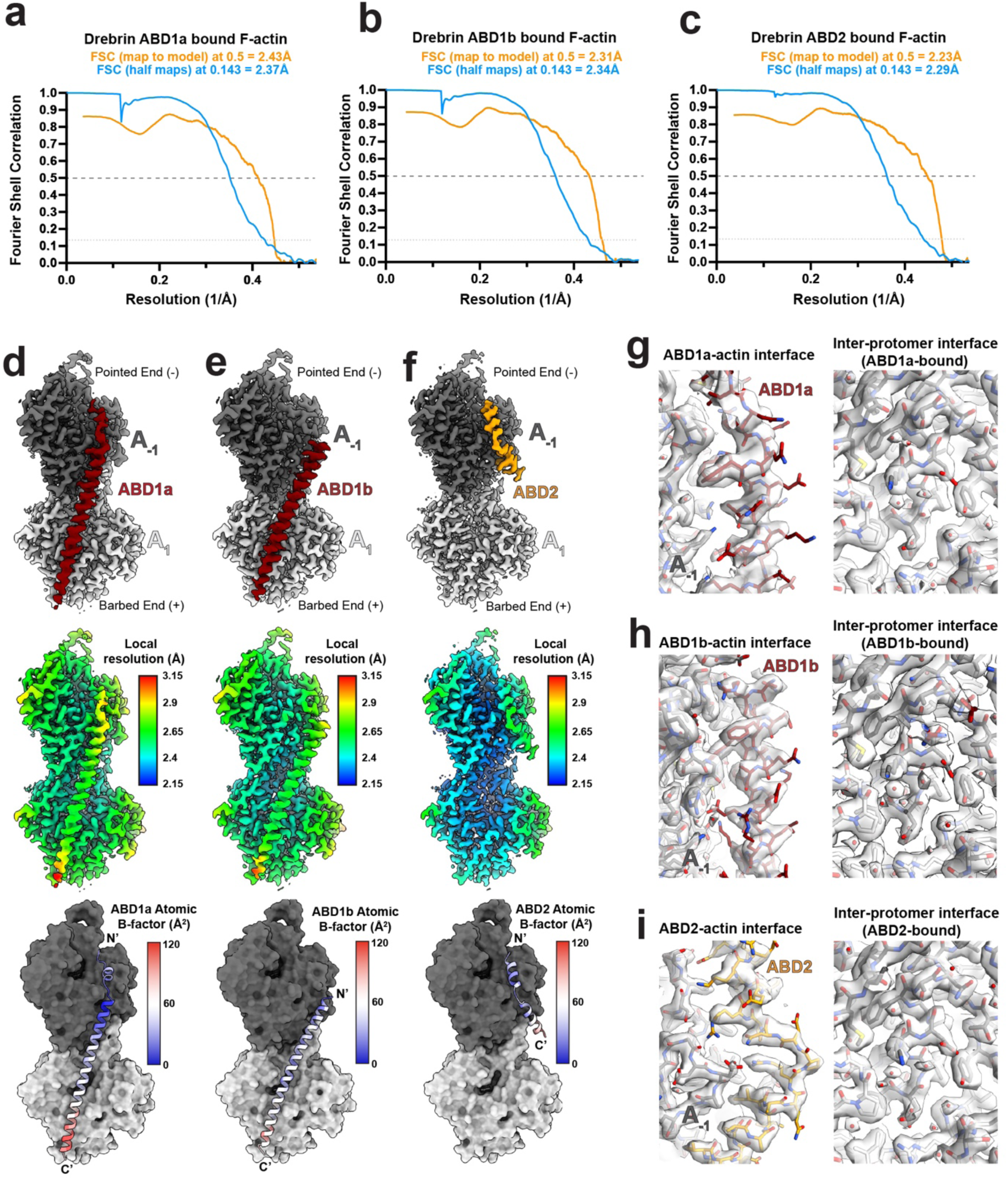
Resolution estimates for ABD1 and ABD2 reconstructions from the drebrin-E^135–355^-decorated F-actin dataset. Gold-standard Fourier shell correlation (FSC) curves between independent half-maps (corrected after noise-substitution tests^104^, calculated in RELIONv5^99^) and map-model FSC curves are shown for the focussed reconstructions of a) ABD1a, b) ABD1b and c) ABD2-decorated F-actin from our drebrin-E^135–355^ dataset. Resolution estimates are shown above, based on the 0.143 cutoff criterion for half-map FSCs and the 0.5 cutoff criterion for map-model FSCs. Focussed reconstructions are shown of d) ABD1a, e) ABD1b and f) ABD2-decorated F-actin from the drebrin-E^135–355^ dataset, coloured by domain (top panels) or the same view filtered and coloured by local resolution using RELIONv5^99^ local resolution estimation (middle panels). For each reconstruction, the respective fitted model is shown in the bottom panels, with F-actin displayed as a surface and the drebrin ABD as a cartoon representation coloured by atomic B-factor. Example densities and fitted models for the drebrin-F-actin interface (left panels) and the central inter-protomer interface in F-actin (right panels) are shown for our g) ABD1a, h) ABD1b and i) ABD2-decorated F-actin reconstructions. All densities are sharpened to local resolutions determined using RELIONv5^99^ and displayed at an RMS deviation threshold of 12.

**Extended Data Fig. 5.**
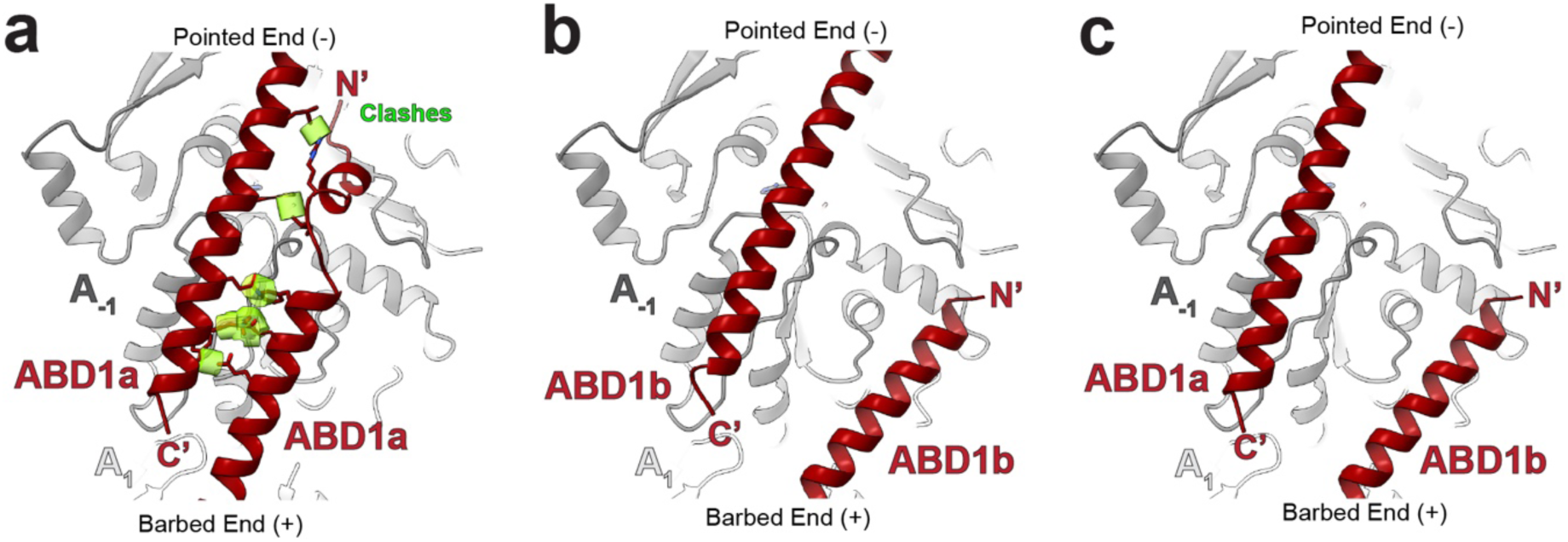
Definition of F-actin binding stoichiometry for ABD1 conformations. a) Our ABD1a F-actin model is superimposed via protomer A_1_ onto another ABD1a F-actin model with its protomer A_-1_ as a reference. b) Our ABD1b F-actin model is superimposed via protomer A_1_ onto another ABD1b F-actin model with its protomer A_-1_ as a reference. c) Our ABD1a F-actin model is superimposed via protomer A_1_ onto an ABD1b F-actin model with protomer A_-1_ as a reference. Clashes between ABD1s after this superimposition were detected with default settings using ChimeraX’s ‘clashes’ tool^103^) and displayed as transparent green density.

**Extended Data Fig. 6.**
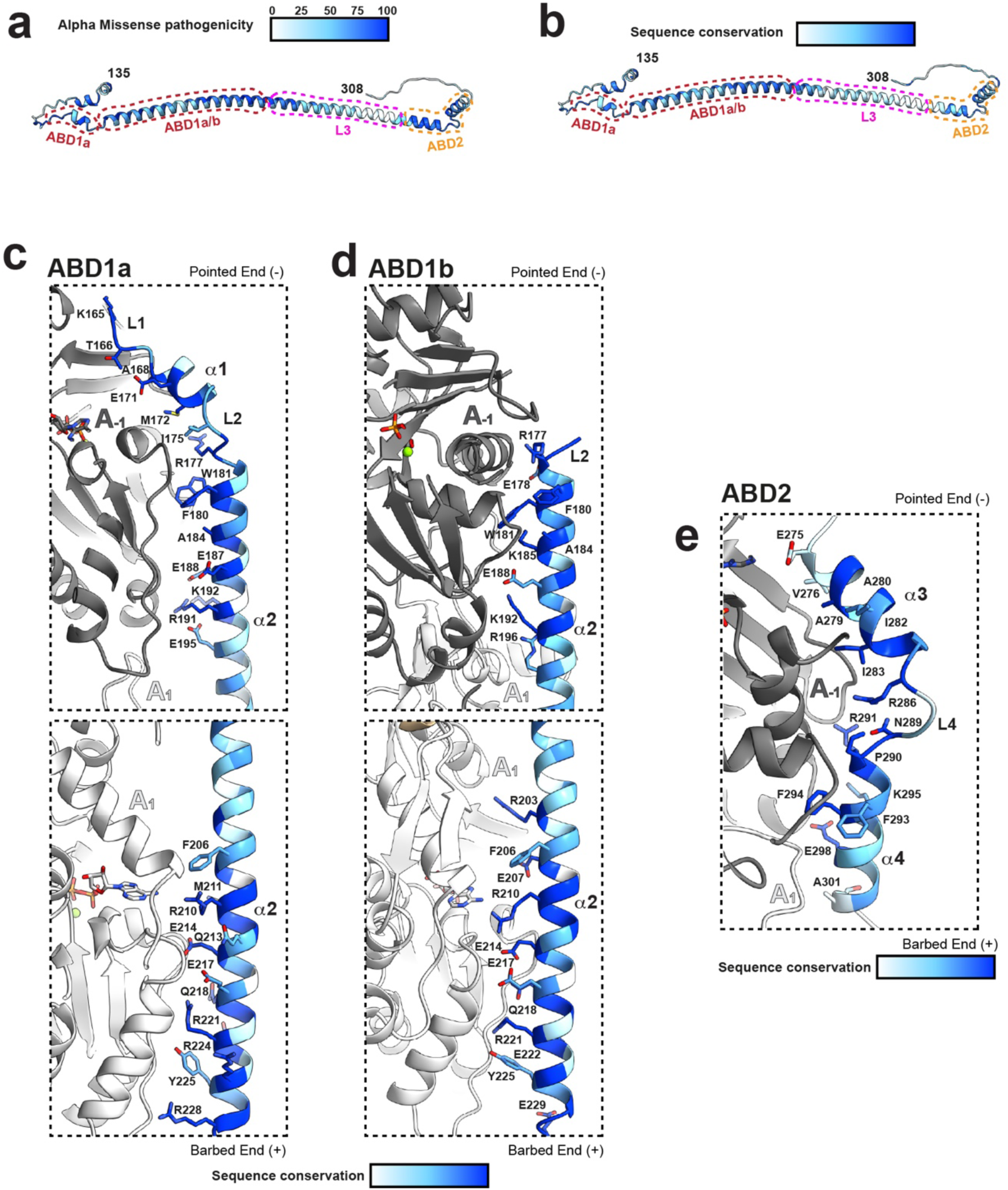
F-actin-interacting residues in Drebrin ABDs are highly conserved and important for drebrin function. The Alphafold3 predicted structure of drebrin-E with residues 135-308 coloured by a) AlphaMissense^90^ pathogenicity scores or b) sequence conservation calculated from the 114 species sequence alignment, with ABDs and L3 indicated with dashed lines. Our cryo-EM-derived drebrin-E-bound F-actin models are shown for c) ABD1a, d) ABD1b and e) ABD2, with drebrin-E coloured by conservation from the 114 species alignment and key F-actin interacting side chains shown/labelled. ABD1 models are broken into two images to accommodate the best views.

**Extended Data Fig. 7.**
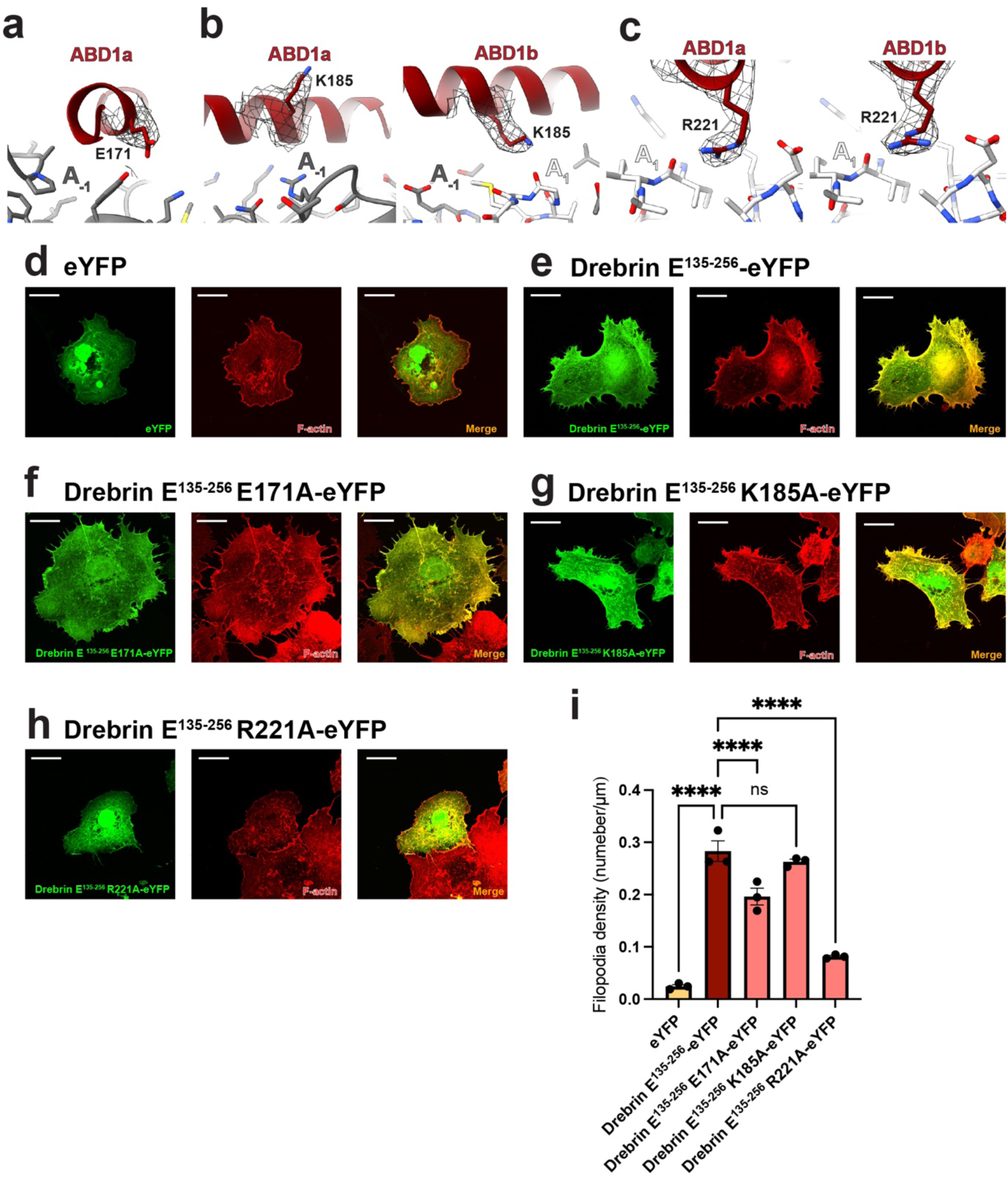
Mutations to F-actin interacting residues reduces ABD1’s ability to induce filopodia in COS7 cells. Zoomed images of our ABD1 F-actin models focussed on drebrin-E amino acids a) E171 (ABD1a; residue is in helix-α1 which is only ordered/interacting with F-actin in the ABD1a conformation), b) K185 (left panel; ABD1a, right panel ABD1b) or c) R221 (left panel; ABD1a, right panel ABD1b). Cryo-EM density for each side chain is shown in mesh (sharpened with DeepEMhancer^106^). Representative examples of COS7 cells transfected with d) eYFP, e) drebrin-E^135–256^-eYFP, f) drebrin-E^135–256^-E171A-eYFP, g) drebrin-E^135–256^-K185A-eYFP and h) drebrin-E^135–256^-R221A-eYFP. YFP and phalloidin-AF568 fluorescence channels are shown. Scale bars = 30µm. i) Quantification of filopodia density (number of filopodia per unit length of cell perimeter) in COS7 cells. For each condition, 3 independent transfections were performed then measurements taken from 10 cells from each transfection. Each data point represents the mean of 10 cells from a single independent transfection. Error bars represent ± standard error of the mean (SEM) for each condition. Ns = not significant, **** = p<0.0001, 2-way ANOVA.

**Extended Data Fig. 8.**
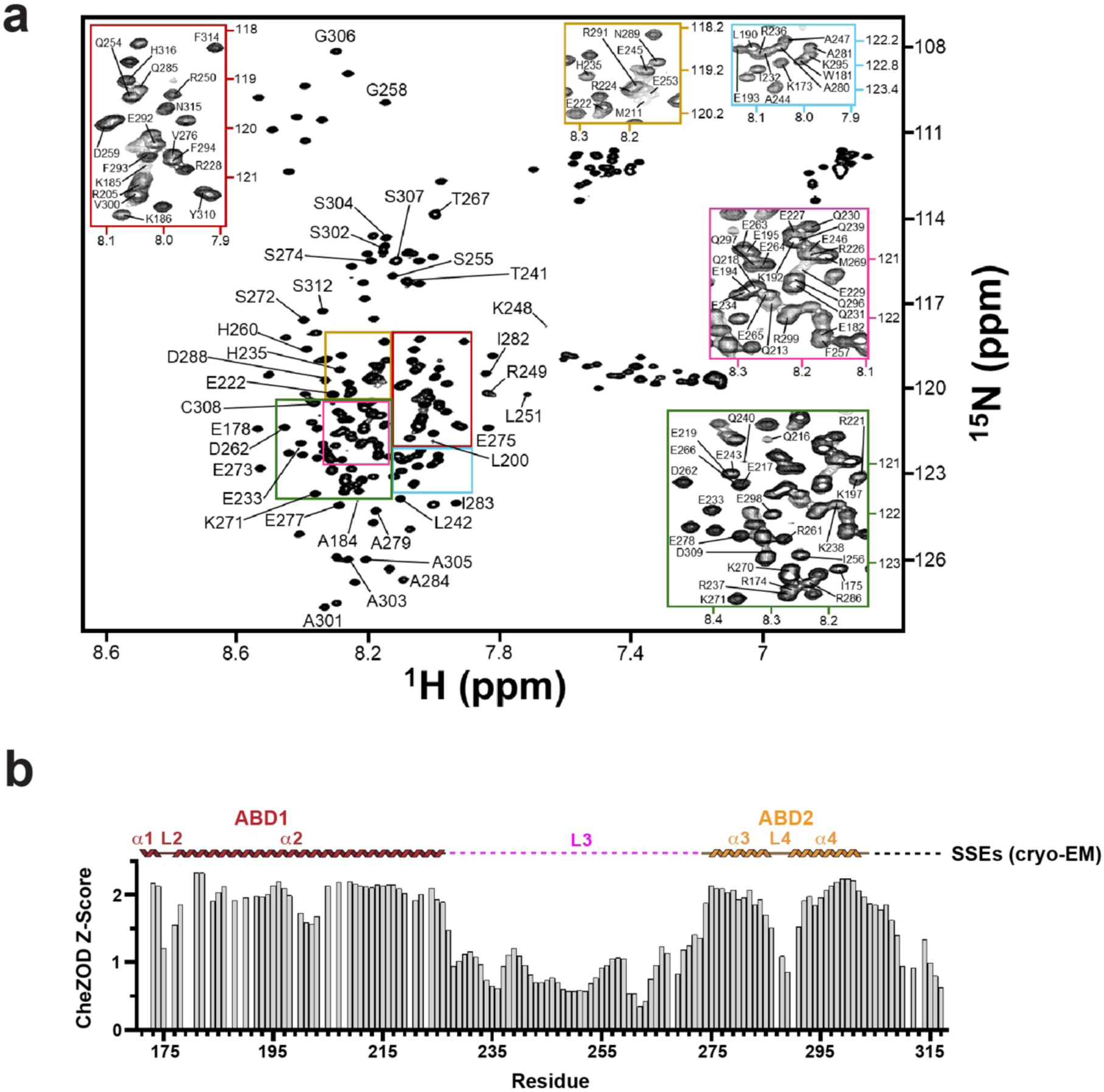
NMR of drebrin-E^135–355^. a) Assigned ^1^H-^15^N HSQC NMR spectrum of ^15^N-drebrin-E^135–355^. The coloured insets are zoomed in crowded regions of the spectrum. The spectrum was assigned using previous NMR studies of drebrin fragments spanning residues 173–238 and 233–317^91,92^ giving 94% coverage of the sequence spanning 173-317. b) cheZOD^70^ secondary structure propensity (SSP) scores for drebrin-E residues 173-317 calculated from the assigned ^1^H-^15^N HSQC NMR spectrum of ^15^N-drebrin-E^135–355^. Higher scores correspond to greater secondary structure propensity, whereas CheZOD values of 0 indicate unassigned residues. The experimentally observed secondary structure elements (SSEs) derived from our cryo-EM data are displayed above the corresponding residue numbers.

**Extended Data Fig. 9.**
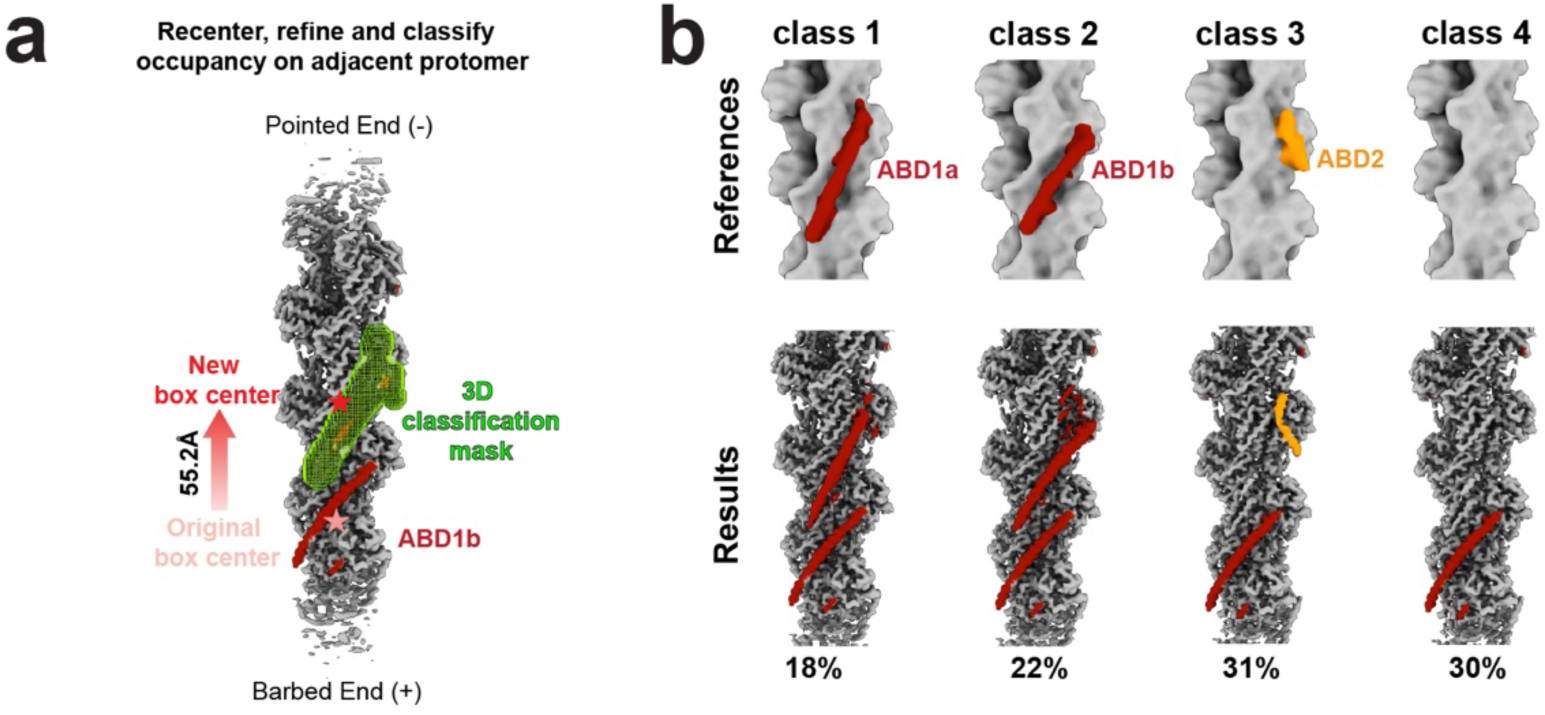
There is no clear spatial relationship between ABD1b and other ABDs on F-actin. a) Schematic showing classification strategy to assess occupancy of ABDs at F-actin substrate sites adjacent to ABD1b. Density from the refined reconstruction is shown after re-extracting/recentring 55.2 Å towards the pointed end (new box centre, red) of the original reconstructed ABD1b-bound class (original box centre, faded red). The focussed 3D classification mask for ABD occupancy is shown in green. b) Corresponding supervised 3D classification class references and results, derived from the strategy indicated in panel a. Density is coloured according to molecular identity (F-actin = grey, ABD1 = maroon and ABD2 = orange) and class occupancy % is shown below the results.

**Extended Data Fig. 10.**
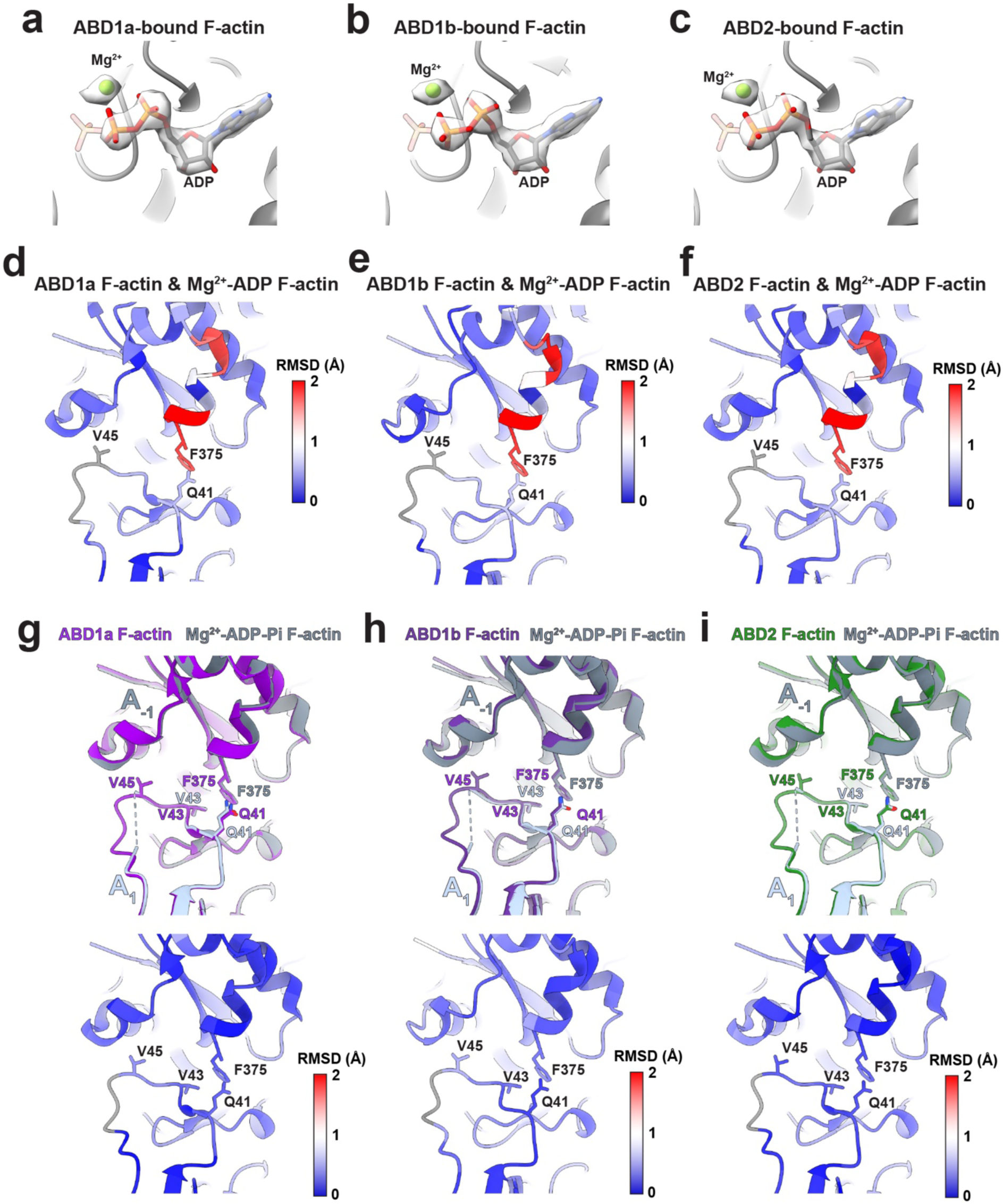
Drebin-E ABDs stabilise a D-loop and C-terminus conformation seen in Mg^2+^-ADP-Pi-bound F-actin. Zoomed density and model for Mg^2+^-ADP associated with the a) ABD1a, b) ABD1b and c) ABD2-bound F-actin reconstructions. The theoretical position of the γ-phosphate if ATP was bound is shown as a faded model. Cα Root Mean Square Deviations (RMSDs) corresponding to the superimpositions of Mg^2+^-ADP F-actin alone (PDB: 8A2T^67^) with d) drebrin ABD1a-bound, e) drebrin ABD1b-bound and f) drebrin ABD2-bound Mg^2+^-ADP F-actin shown in Fig. 5d-f (view focussed on the D-loop and C-terminus). View focussed on the F-actin D-loop and C-terminus after superimpositions of our g) ABD1a Mg^2+^-ADP F-actin, h) ABD1b Mg^2+^-ADP F-actin and i) ABD2 Mg^2+^-ADP F-actin models with the model of Mg^2+^-ADP-Pi-bound F-actin alone (PDB: 8a2s^67^). Cα RMSDs corresponding to the superimpositions are shown below.

## Notes

### Competing Interest Statement

The authors have declared no competing interest.

